# Unsupervised generative AI discovers pan-leukocyte dysregulated pathways in single-cell lupus data

**DOI:** 10.1101/2024.03.02.583143

**Authors:** Hamid Bolouri, Karen Cerosaletti, Adam Lacy-Hulbert

## Abstract

Comparisons of single-cell RNA-sequencing (scRNA-seq) data from healthy and diseased tissues are widely used to identify cell-type-specific dysregulated genes, pathways, and processes. Accordingly, many sophisticated methods have been developed for this purpose. However, such tools generally require considerable user expertise for optimal performance. Here, we show that unsupervised application of a linearly-decoded Variational Auto Encoder (a generative AI model) to scRNA-seq data recapitulates and extends findings from a seminal recent lupus study and leads to new insights. Thanks to existing software libraries, our approach is straightforward to implement, computationally efficient, methodologically robust, and produces consistent and reproducible results.

## lntroduction

A wide variety of computational methods have been developed to infer molecular processes dysregulated in diseased cells using single-cell RNA-sequencing (scRNA-seq) data. Common approaches range from direct healthy-disease comparisons such as differential-expression ^1^ and differential abundance analysis ^2^, to inference of multi-gene health-to-disease expression changes via trajectory analysis ^3,4^, RNA-velocity ^5^, signaling analysis ^6^, and gene-co-expression module detection ^7,8^. Simple comparisons typically suffer from limited power. Multi-gene inference-based methods offer deeper, more robust insights but typically involve expert tuning of various ad-hoc analysis steps as well as method configuration choices (see for example ^9,10^). As scRNA-seq technology becomes cheaper and more high-throughput, there is a need for methods that allow robust, unsupervised, and easy-to-automate multi-gene case-control comparisons.

Variational Auto Encoders (VAEs) ^11^ combine the learning power of generative AI models with the computational efficiencies of latent variable models. However, the latent representations of VAEs, while efficient, are nonlinear and therefore not easily interpretable. To overcome this issue, Svensson et al recently proposed a modified VAE ^12^ in which the encoder is nonlinear but the decoder is linear. Such linearly decoded VAEs (ldVAEs) have the advantage that their latent space dimensions are each a linear weighted sum approximation of the input variables (e.g. gene expression levels).

Here, we show that when applied to scRNA-seq data from patients and healthy-donor (HD) controls, ldVAEs reliably and consistently identify disease associated genes and their relative importance. In addition, we show that ldVAEs require no supervision/expert-guidance and their findings are robust to network configuration and runtime parameters. Using Systemic Lupus Erythematosus (SLE) as an exemplar of a highly complex disease that dysregulates multiple cell types, we demonstrate the usefulness of ldVAE models by showing that ldVAE results recapitulate and extend published findings and enable novel insights about SLE pathogenesis.

SLE is a highly heterogeneous autoimmune disease ^13^ that disproportionately impacts young women of color^14^. SLE epitomizes the challenges facing complex-disease case-control studies ^15^ in that it impacts diverse organs, and its effects vary from person to person and over time ^16,17^. Moreover, subsets of B cells ^18^, T cells ^19^, monocytes ^19^, and dendritic cells (DCs) ^20^ have all been shown to be dysregulated in SLE. The involvement of many different genetic and environmental factors, organs, cell types, and molecular processes have hindered the development of curative and targeted SLE treatments ^21^. In particular, it is not clear whether common mechanisms drive flares across all SLE patients.

In recent years, two broad-brush theories of SLE-pathogenesis have gained increasing acceptance. First, immune dysregulation in SLE is thought to be due to a self-reinforcing feedback loop between inflammation and high levels of cell/organelle stress, damage, and death that overwhelms the cellular debris-clearance apparatus ^22,23^. Second, diverse drivers of SLE pathogenesis may impact a small number of key disease drivers. In particular, the X-chromosome has emerged as a potential common target of diverse factors driving SLE pathogenesis. Consistent with this hypothesis, auto-antibodies to the X-chromosome inactivating gene XIST ^24^ and escape from X-Chromosome Inactivation (XCI) ^25^ have been implicated in SLE pathogenesis and may explain the predominance of women in SLE ^25,26^. Here, our findings identify a potential link between the inflammatory-feedback and X-chromosome hypotheses.

In what follows, we first present the results of applying ldVAEs to blood immune cell scRNA-seq data from HD and lupus flare patients. We then evaluate the extent to which unsupervised ldVAE findings confirm and extend findings from previous studies. Finally, to verify and pinpoint potential causal mechanisms underlying the ldVAE findings, we perform additional network-based integrative analysis of heterogeneous data from seven single-cell and sorted-cell datasets from SLE and healthy blood donors (summarized in Supplementary Figure 1). Results from this extended analysis support the ldVAE findings and provide a novel hypothesis connecting dysregulated debris-processing with IFNα/β signaling and XCI escape.

**Figure 1.**
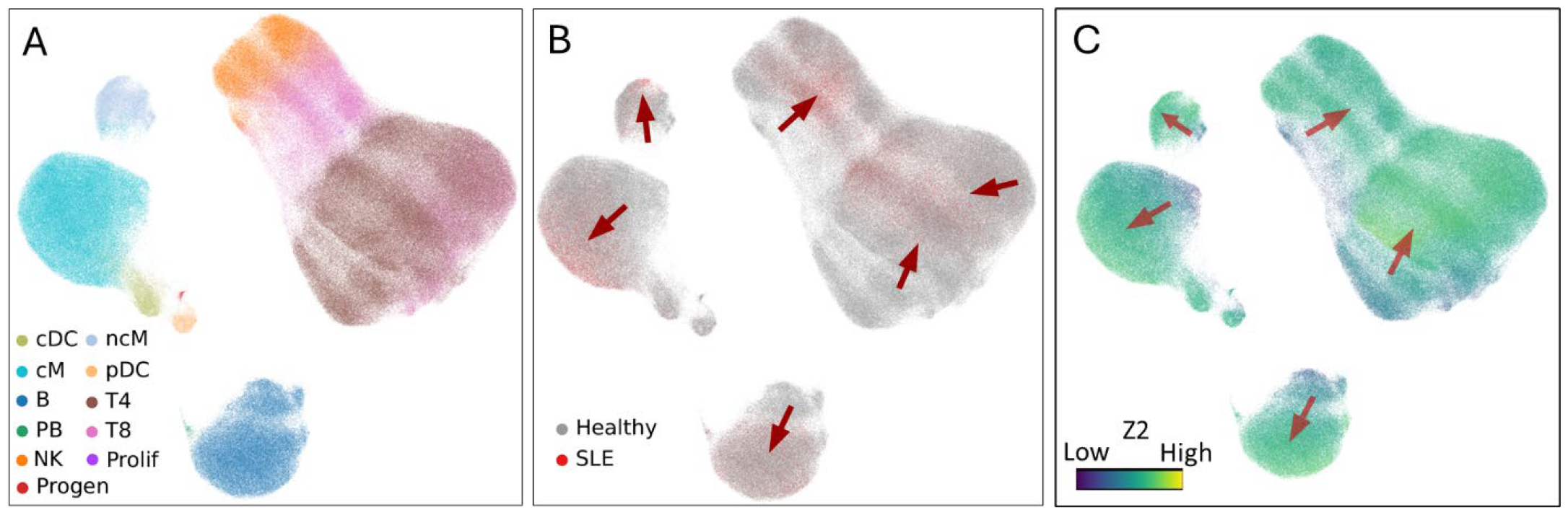
Characteristics of an example ldVAE model (denoted L1H128Z8) trained on scRNA-seq data for 55,120 SLE-Flare and 486,418 Healthy Donor PBMCs. (**A**) Two-dimensional UMAP visualization of the latent space of the model colored by the original cell labels assigned by Perez et al. (**B**) Same UMAP as in (A) here colored to show the distributions of cells from Healthy Donors and SLE patients. Arrows highlight gradients of increasing SLE cell density. (**C**) The same UMAP as in (A) and (B) colored by the values of latent dimension Z2. Arrows highlight gradients of increasing Z2.

## Results

### Data characteristics and pre-processing

We used large-scale scRNA-seq data from peripheral blood mononuclear cells (PBMC) published in a seminal recent study by Perez et al ^27^ to train a ldVAE. A landmark earlier study in a smaller SLE cohort ^28^ found that – irrespective of age and other factors – SLE patients undergoing a flare exhibit a strong interferon-response signature across all major peripheral blood immune populations. Accordingly, we focused on the subset of SLE patients undergoing a flare. To exclude treatment effects, we selected 55,120 PBMCs collected from 19 SLE patients at the onset of a flare. As controls, we used scRNA-seq data from the same study for 486,418 PBMCs from 99 HDs. For each training run, 10% of the data was randomly excluded from training and used to test and validate the model.

### ldVAEs consistently identify genes up-regulated in SLE

We trained a total of 10 ldVAEs with varying numbers of hidden layers and hidden layer nodes to compress the training data into a range of four to 15 latent dimensions. As shown in **Figure 1A** and Supplementary Fig. 2, all 10 models generated low-dimensional latent spaces in which cells of the same type cluster together. This finding confirms that our ldVAE models capture key biological properties of SLE and HD single-cell transcriptomes.

**Figure 2.**
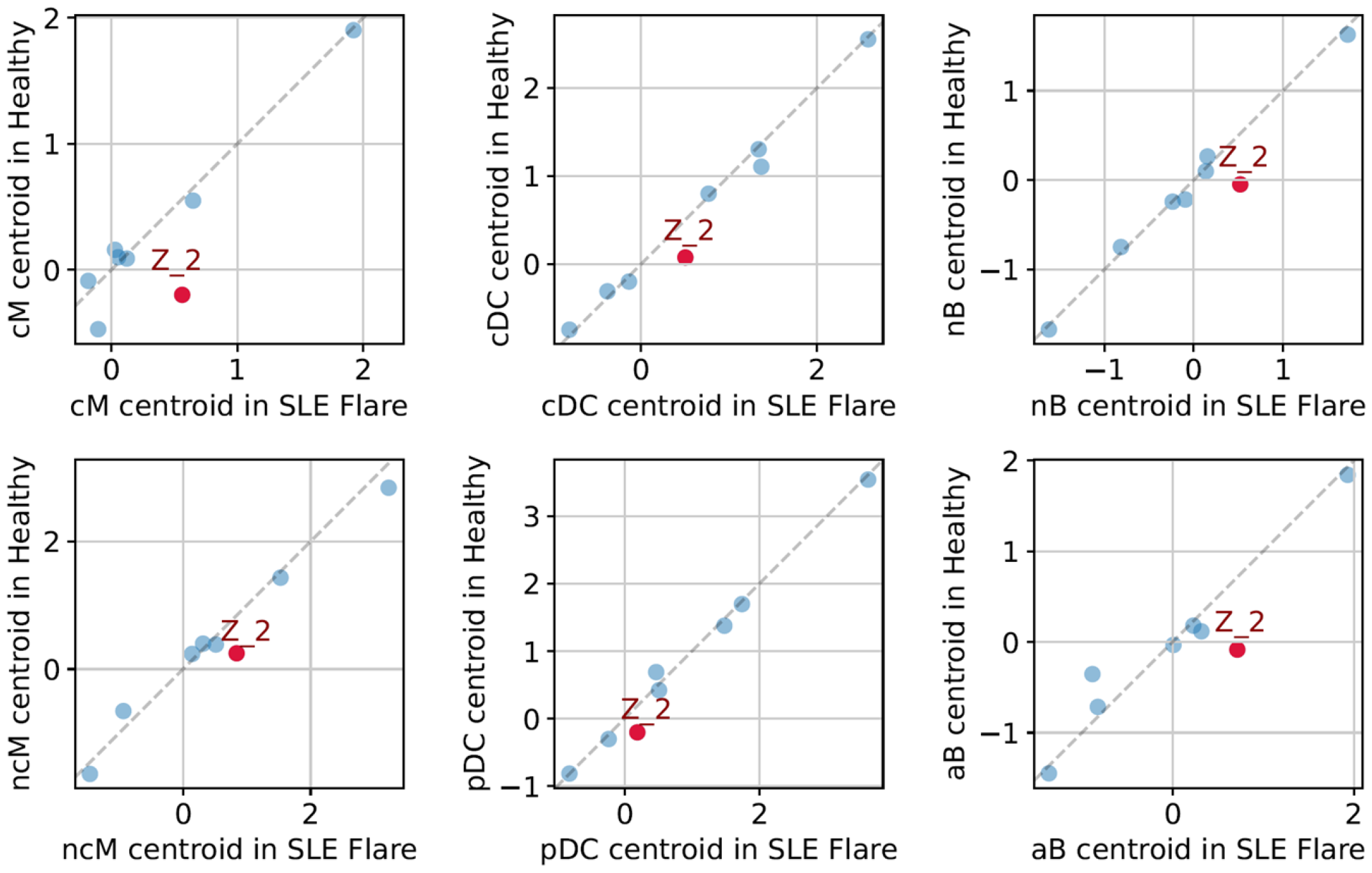
SLE leukocytes have higher average levels of latent dimension Z2 in model L1H128Z8 (see Fig. 1). Each panel compares healthy versus SLE mean Z2 values for a different immune population identified by Perez et al: cM: classical monocytes, ncM: non-classical monocytes, cDC: conventional dendritic cells, pDC: plasmacytoid dendritic cells, nB: naïve B-cells, aB: atypical B-cells. Dashed lines mark locus of equal values for healthy and SLE cells. Each plot-point represents one of the model’s eight latent dimensions. In all size immune populations, average Z2 is higher in SLE compared to hD.

While healthy and SLE cells of the same type cluster together in ldVAE latent space, we observed a distinct healthy-to-disease gradient within each cell-type cluster (**Figure 1B** and Supplementary Fig. 2), suggesting the ldVAE models capture key transcriptomic differences between HD and SLE immune cells. Remarkably, we find that the HD-to-SLE gradients are mirrored by changes in specific latent dimensions (see in **Figure 1C** and Supplementary Figure 2).

To identify cell-type-specific HD to SLE expression changes, for each cohort, we calculated the mean value (centroid) of each immune cell population for each latent dimension. Comparing per-cell-type HD versus SLE latent-dimension centroids, we found that in seven of our ten ldVAE models, the most discriminative latent dimension was the same across multiple myeloid and lymphoid cell types (**Figure 2** and Supplementary Fig. 2).

By design, each ldVAE latent dimension is a linear weighted sum of the input gene expression values, and the weight associated with each gene determines the relative contribution of that gene to changes along the respective latent dimension. Focusing on genes that vary most (i.e. have the largest absolute weight values) along the identified healthy-to-SLE latent dimension (see example in **Figure 3**), we found that in all seven models the top 25 genes with the greatest positive effect size (weight) are highly enriched for three biological processes: Interferon signaling, cellular debris clearance, and immune dysregulation (including autoimmunity and loss of immune tolerance, see **Figure 4A, B** and Supplementary Fig. 3). In contrast, genes down-regulated in SLE did not show a consistent enrichment across models and are not discussed further.

**Figure 3.**
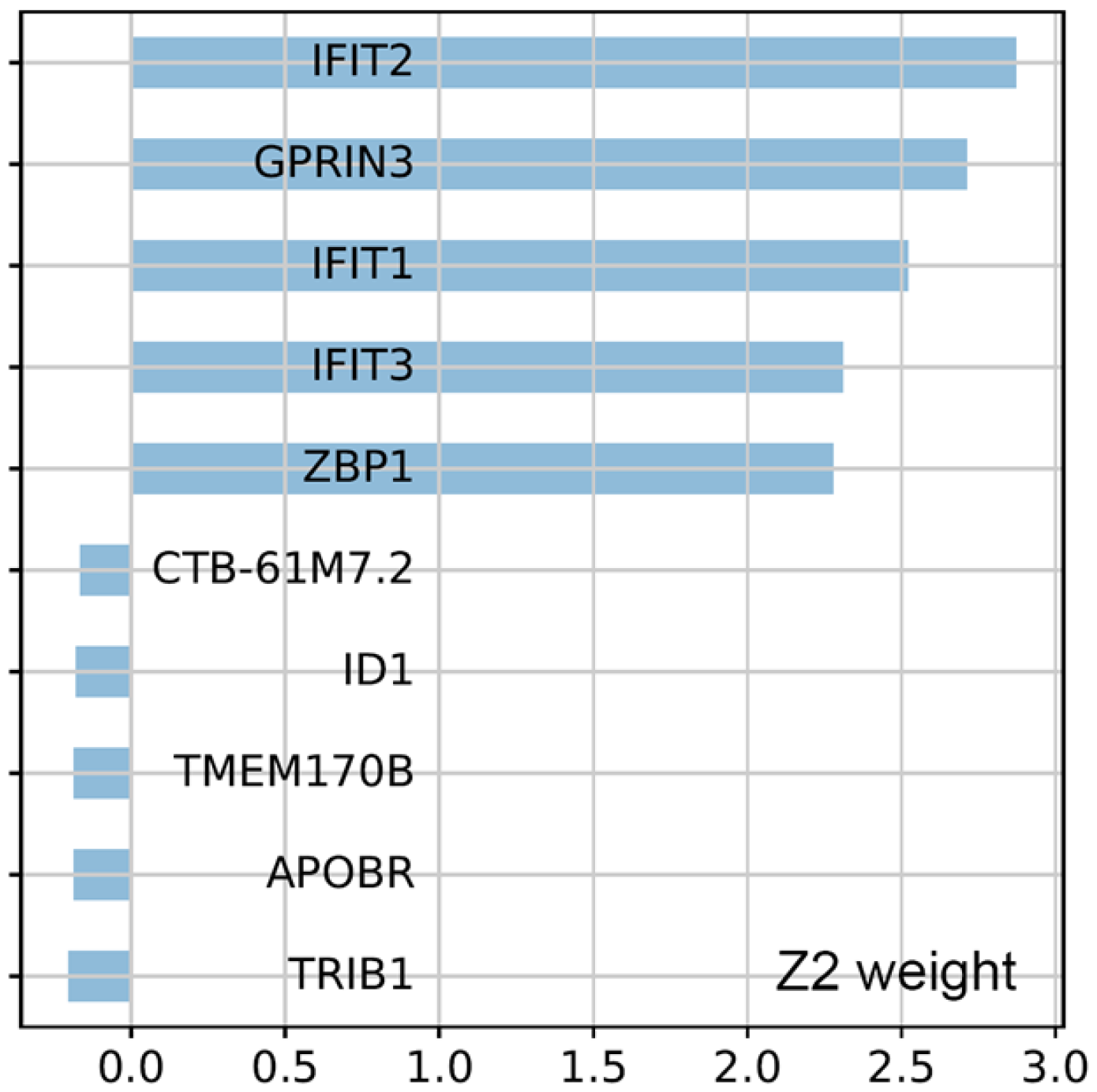
Top-5 genes with the highest positive and negative weights in latent dimension Z2 of the ldVAE model in figures 1 and 2. As shown in figs. 1C and 2, Z2 is increased in SLE vs. HD. As shown here, increases in Z2 are primarily driven by higher interferon signaling genes (IFIT 1/2/3), debris-clearance associated genes (ZBP1) and autoimmunity (GPRIN3). For additional genes, see Fig. 4A and Supplementary Fig. 3.

**Figure 4.**
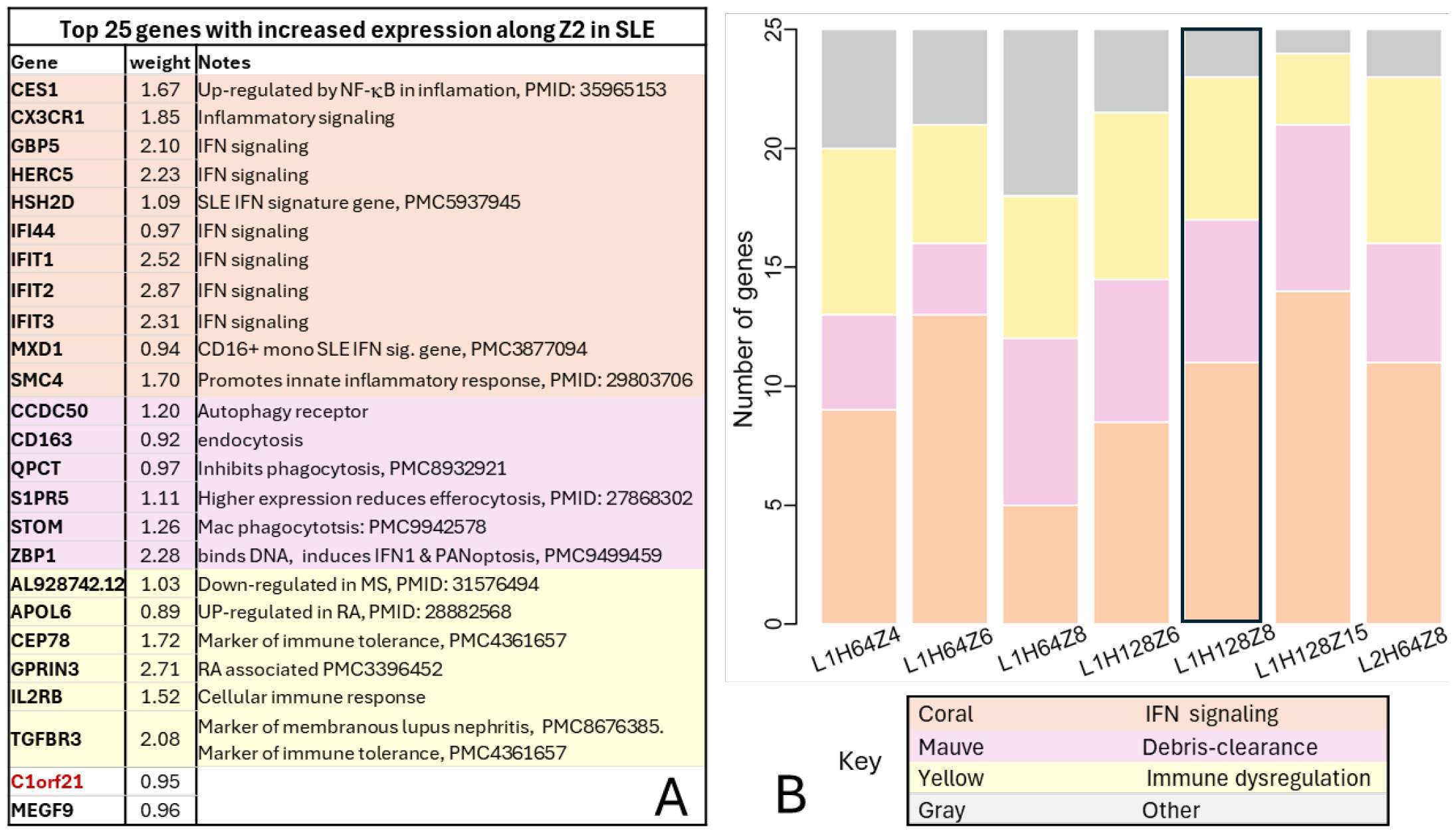
SLE-specific ldVAE latent dimensions are highly enriched for IFN signaling, debris-clearance, and immune dysregulation genes. (**A**) Top 25 genes increasing with higher Z2 weights for the model in figs. 1 & 2. (**B**) Frequencies of genes associated with the same processes as in (A) in seven independent ldVAE models. Model L1H128Z8 (panel (A) and figs 1 and 2) is highlighted. Model names describe their network configuration in the format ‘LxHxZx’, where L stands for number of layers in the encoder network, H stands for the number of hidden units, Z stands for the number of latent dimensions, and x is a number. Thus, for example, ldVAE L1H64Z4 has a decoder with 1 hidden layer, 64 hidden neurons, and 4 latent dimensions. All models were trained identically (see Methods).

### Multiple sources of evidence support novel ldVAE findings

Closer inspection of the top-25 SLE-associated genes in the seven ldVAE models revealed that the same genes are often implicated by differently-configured and independently-trained ldVAE models. Indeed, the 175 genes identified by the seven models correspond to 90 unique genes. The gene/protein interaction databases geneMANIA ^29^ and stringDB ^30^ revealed that 86 of these 90 genes have previously been reported to either be co-expressed or to interact physically or functionally (Supplementary Fig. 4). These findings suggest that unsupervised ldVAE training robustly captures biologically relevant disease characteristics in case-control scRNA-seq data.

Given the finding that nearly all the SLE-up-regulated genes identified by ldVAEs have previously been reported to be co-expressed, we hypothesized that many of the same genes will also be found to be co-expressed in healthy PBMCs, which comprised ∼90% of our dataset. To test this hypothesis, we performed transcriptome-wide correlation analysis in HD cells and used label-randomization to select gene-gene correlations with r > 0.5 and empirical p-values < 0.01 (for details see Methods). As illustrated in Supplementary Fig. 5, a large majority of the 90 SLE-upregulated genes identified by ldVAE form fully-connected co-expression networks in HD macrophages, DCs, and B-cells.

**Figure 5.**
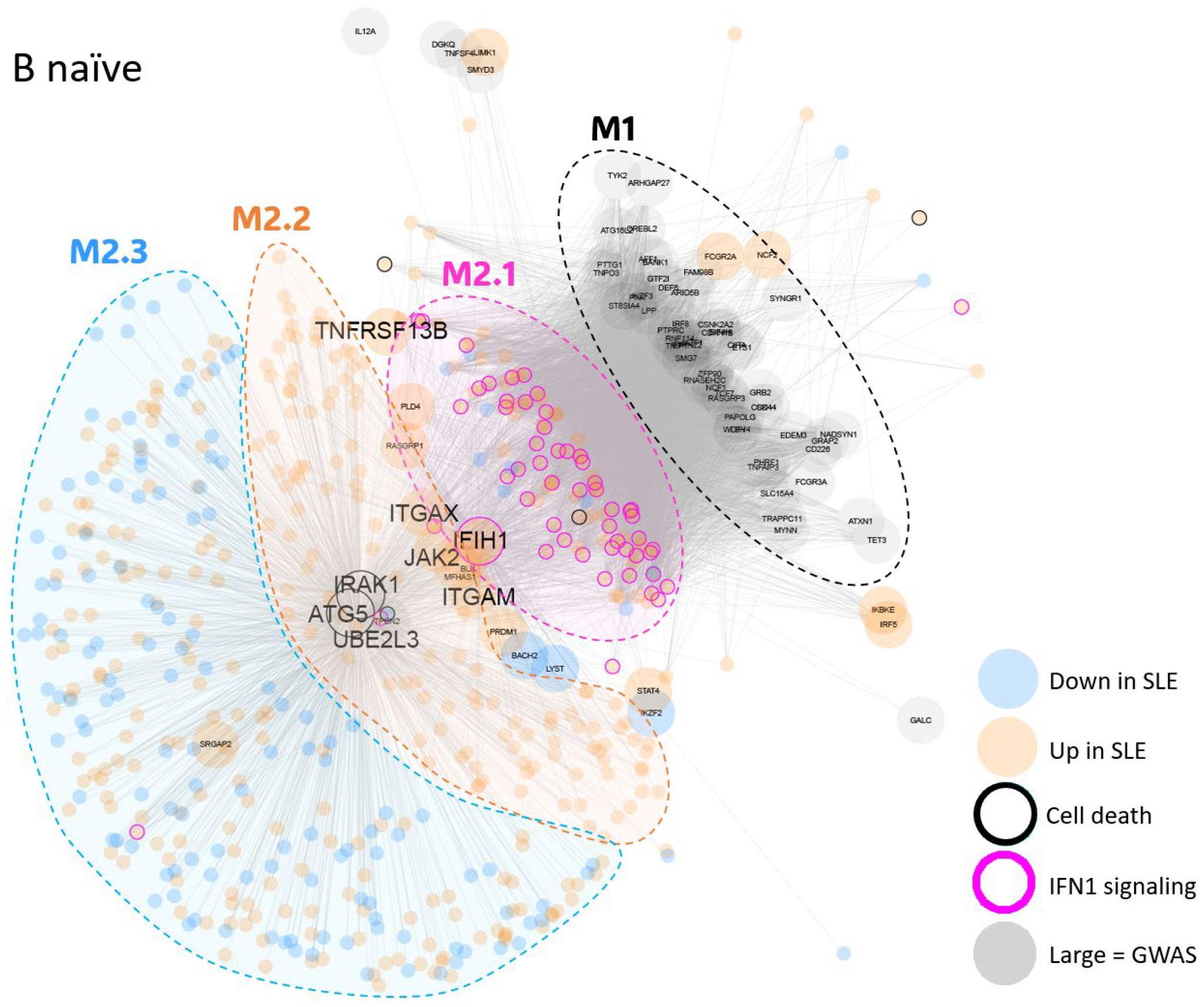
Manually derived SLE-associated gene co-expression network in naïve blood B cells. Nodes represent genes. Edges indicate highly correlated expression in healthy donors. Large nodes represent genes with previously published risk alleles for lupus. Colored nodes are genes differentially expressed between lupus-flare and healthy cells. Magenta and black rings around nodes indicate genes associated type 1 interferon signaling and cell-death/debris clearance. Dashed ovals indicate distinct groups of genes in the network. The network divides into two broad-brush co-expression modules. M1 at top-right is mostly made up of SLE-risk genes that are not differentially expressed in SLE-flare versus healthy cells, but whose expression is nonetheless correlated with type1 interferon signaling genes. Module 2 is centered around three hub genes: ATG5, IRAK1, and UBE2L3. All three of these genes are SLE associated genes involved in cellular debris processing. Module M2 divides into 3 sub-groups: (M2.1) comprises up-regulated IFN1 signaling genes whose expression in HD is highly correlated with diverse SLE-risk genes and also with IFIH1. (M2.2) is made up of other genes up-regulated in SLE and highly correlated with IFIH1. (M2.3) comprises genes highly correlated with ATG5/IRAK1/UBE2L3, but not tightly correlated with IFIH1. GWAS indicates SLE-risk genes identified by Genome-Wide Association Studies.

### ldVAE findings agree-with and extend differential expression analysis

We find that there is significant agreement between ldVAE findings and differentially-expressed genes (DEGs). Of 90 SLE-associated genes discovered by ldVAE models, 37 are among 418 HD versus SLE DEGs identified in all cell-types (hypergeometric enrichment p-value 7.8E-10, see Supplementary Table 1A and Methods for details). Thirteen of these 37 genes are in the Reactome ^31^ ‘R-HSA-913531’ Interferon Signaling pathway (FDR-adjusted hypergeometric enrichment p-value 7.0E-12), and 11 of the 37 genes are in the Gene Ontology ^32^ biological process ‘GO:0012501’ Programmed Cell Death (FDR-adjusted hypergeometric enrichment p-vale 0.0024).

Importantly, in addition to identifying many DEGs, ldVAE modeling identified many SLE-associated genes that are not differentially expressed but impact the pathways enriched in DEGs. Of the 53 SLE-associated genes identified by ldVAE but not by differential expression analysis, seven are associated with interferon signaling, and 17 are implicated in debris clearance (Supplementary Table 1B). Thus, ldVAE analysis confirms the pan-immune-cell interferon signaling reported by Perez et al ^27^ and Nehar-Belaid et al ^28^, and additionally implicates cellular debris clearance as another pan-immune-cell process dysregulated in SLE flare cases.

### ldVAE findings lead to novel insights about SLE pathogenesis

The SLE-upregulated genes identified by our ldVAE models are strongly enriched for two well-established dysregulated molecular processes in SLE: interferon signaling, and cellular debris processing. However, whereas previous studies have focused on distinct types of dysregulation in monocytes/macrophages, DCs, and B cells, our unbiased data-driven findings suggest that a single conserved gene co-expression module associated with interferon signaling and debris-processing is dysregulated in all these immune populations.

To verify the above hypothesis, we augmented our gene co-expression network with single-cell and sorted-cell PBMC expression data from six additional studies (Supplementary Fig. 1). These data span 1,285 healthy donors from control arms of three SLE studies and three HD-only studies ^33-35^. Per cell-type pairwise Pearson correlations were calculated separately in each dataset, and only gene-gene correlations with r > 0.5 and empirical p-value < 0.01 were retained (see Methods).

To focus on genes with a distinct role in SLE, we further filtered the derived gene co-expression networks to keep only gene-pairs in which at least one gene was either an SLE risk gene ^36^ or reported as a DEG in a specific cell type in PBMC scRNSA-seq studies by Perez et al ^27^, Nehar-Belaid et al ^28^, and two PBMC sorted-cell bulk RNA-seq datasets spanning 203 SLE patients ^37,38^.

As illustrated in **Figure 5** and Supplementary Fig. 6, in all immune populations studied, the derived gene co-expression networks are densely-connected, suggesting that the detected genes are functionally related. Consistent with our ldVAE findings, the co-expression networks are structured very similarly across all cell types, and comprise two distinct sub-components: a cluster dominated by SLE-risk genes that are not DEG (module M1) and a large number of co-expressed interferon-signaling associated DEGs (module M2).

**Figure 6.**
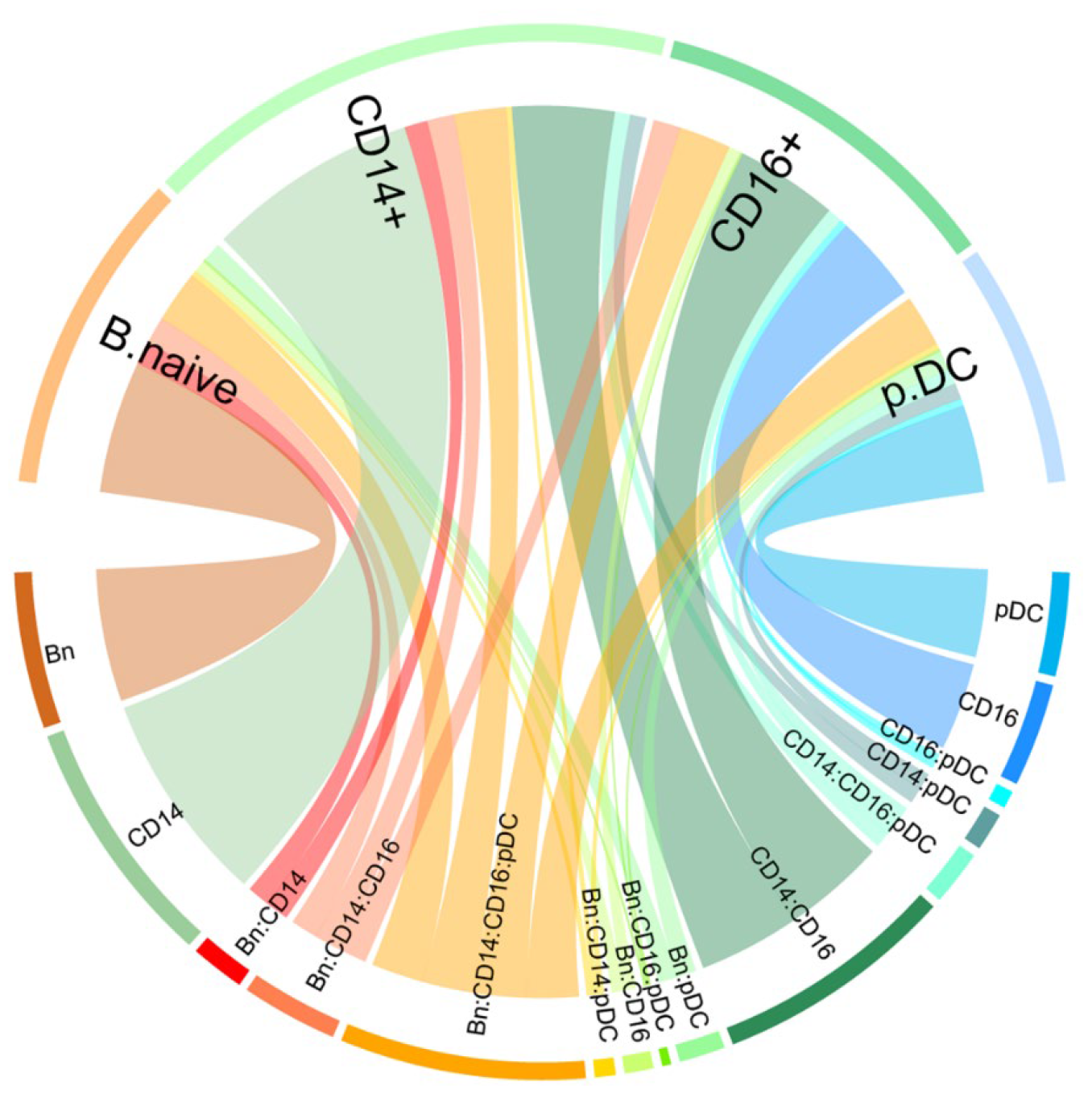
Genes in the M2 module (see Figure 5) are highly conserved across innate and adaptive immune populations. The arcs in the top half represent cells of interest, the arcs in the bottom half represent groups of co-expressed genes. The chords indicate shared co-expression groups. Overlaps are shown for the union of the genes correlated with ATG5, IFIH1, IRAK1, and UBE2L3 across naïve B cells, CD14+ monocytes, CD16+ monocytes, and pDCs. Note that the group of genes shared across all four cell types (labeled “Bn:CD14:CD16:pDC”, colored orange at bottom) is the largest shared group.

Although module M1 genes are associated with increased genetic risk for SLE, very few are differentially expressed compared to HD. Nonetheless, the expression patterns of module M1 genes are highly correlated with the expression levels of a set of Type 1 Interferon activated genes that make up part of module M2 (sub-module M2.1 in **Figure 5**). Module M2 can be divided into three sub-modules. Genes in sub-modules M2.1 and M2.2 are highly correlated with the interferon induced gene IFIH1. In contrast to M2.1 and M2.2 genes, which are up-regulated in SLE, module M2.3 includes both up and down-regulated genes. Nonetheless, genes in M2.3 are also highly enriched for Type 1 Interferon signaling (Supplementary Fig. 7).

**Figure 7.**
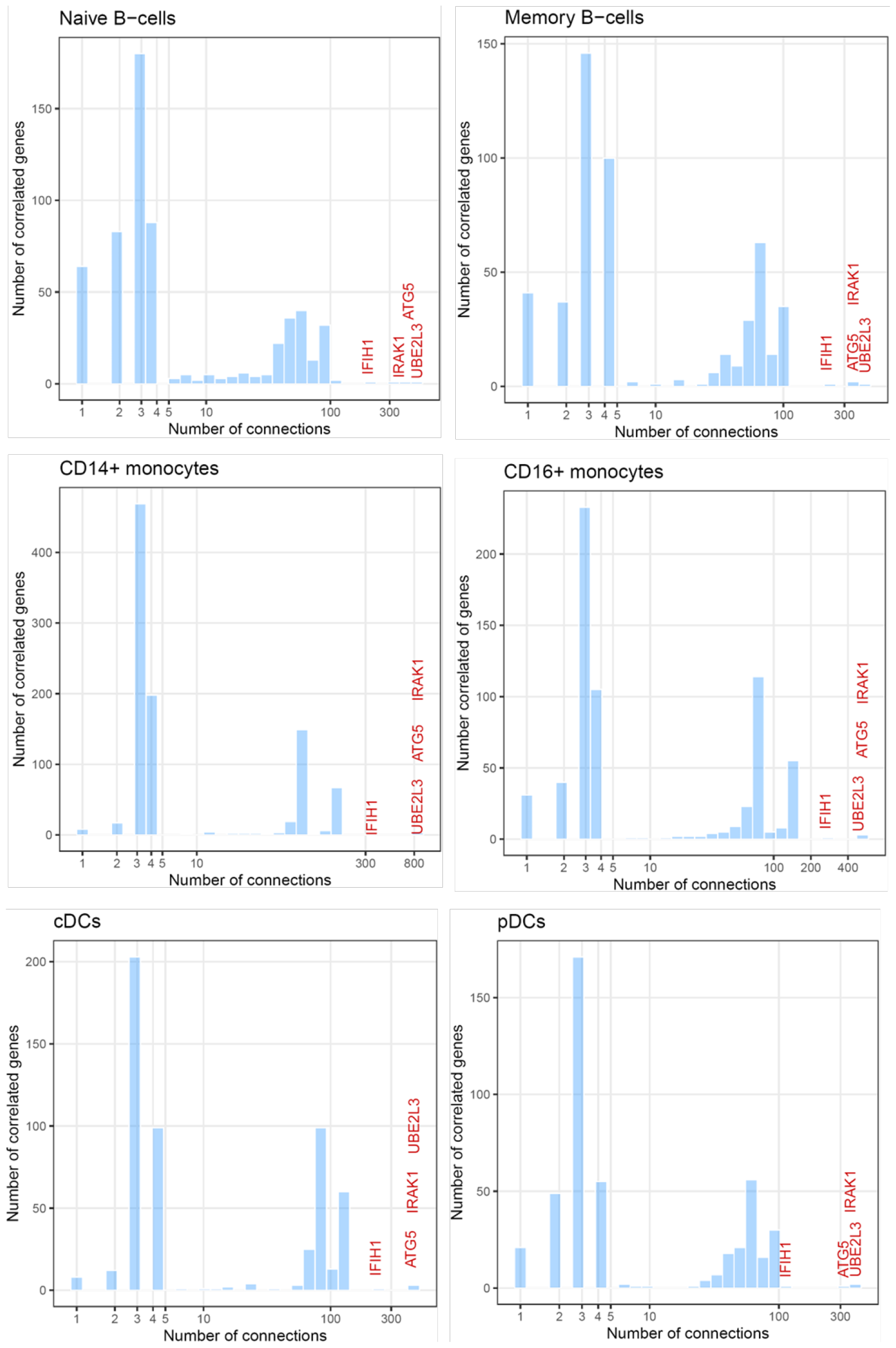
ATG5, IFIH1, IRAK1, and UB2L3 are the most highly connected correlation-network hubs across B cells, monocytes, and DCs. The X-axis shows the number of connected (highly correlated) genes per network node (gene) in each cell type. The Y-axis shows the number of genes with that many connections.

As shown in **Figure 6**, the genes in module M2 are highly conserved across naïve B-cells, CD14+ and CD16+ monocytes, and pDCs. This finding further supports the ldVAE results suggesting that a shared set of interferon-responsive genes are up-regulated in both myeloid and lymphoid cells in SLE versus HD..

Of particular relevance here, nearly all genes in module M2 are connected to three SLE-risk genes with known roles in debris-clearance: ATG5, IRAK1, and UBE2L3. Indeed, across all genes in modules M1 and M2, these three genes are the most-connected network hubs in B cells, monocytes and DCs (**Figure 7** and Supplementary Tables 2-5). The fourth most connected gene across M1 and M2 is IFIH1, which connects M1 and M2. IFIH1 encodes MDA5, an intracellular double-stranded RNA sensor that is involved in both debris-clearance and Type 1 Interferon signaling ^39^. Thus, the four top network hubs in our SLE networks connect debris-clearance and IFNα/β signaling.

It is striking that although module M2 is centered on three debris-clearance genes, it is highly enriched for interferon signaling genes. In addition to being functionally related, the genes correlated with each of the top-4 hub genes are highly conserved across diverse immune cell types (**Figure 8** and Supplementary Fig. 8). Thus, our investigator-driven integrative network analysis confirms the findings of our unsupervised ldVAE models that a common gene-co-expression network centered on debris-clearance and highly enriched for interferon signaling is dysregulated across B cells, monocytes and DCs during SLE flares.

**Figure 8.**
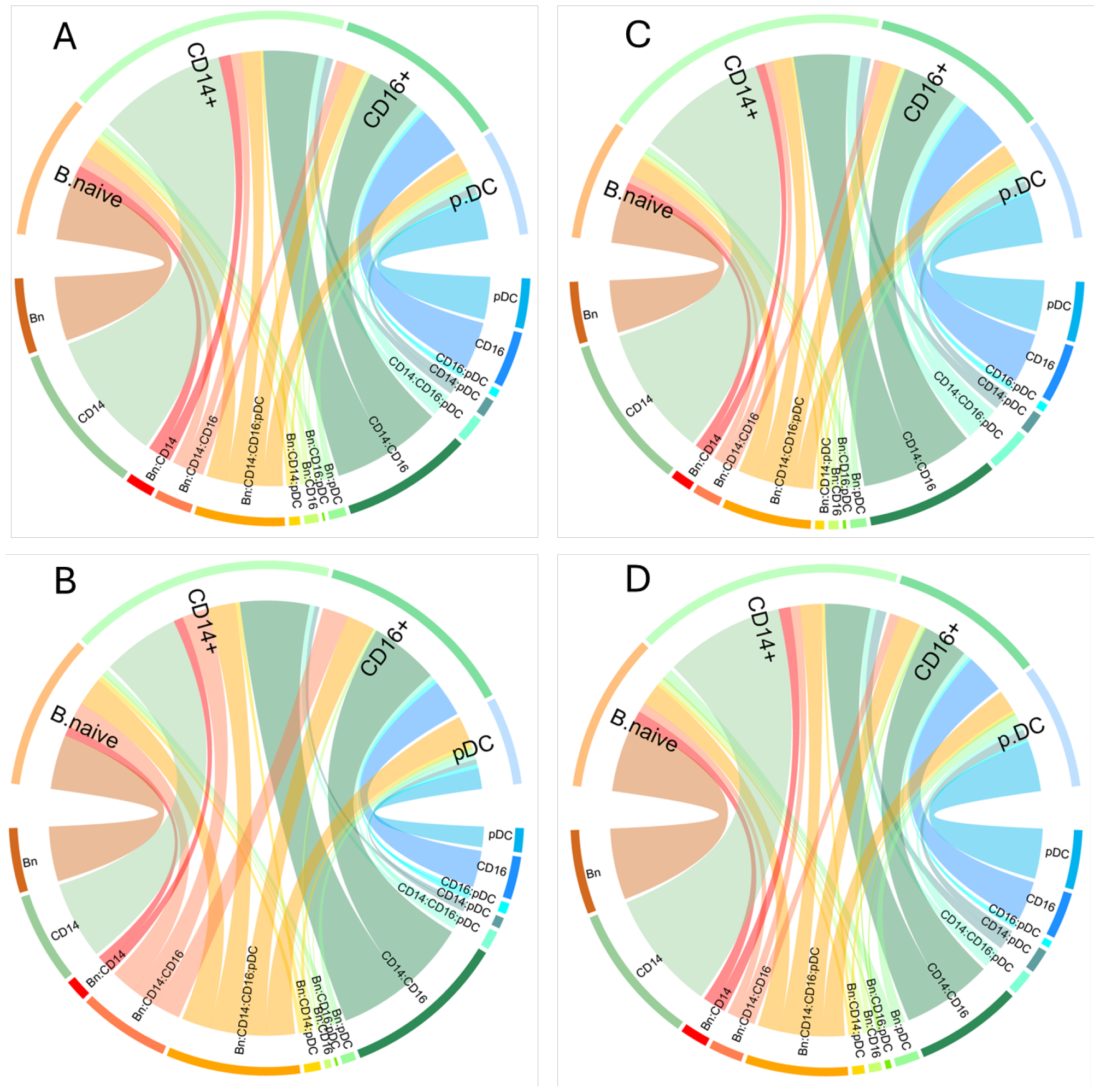
A large proportion of genes correlated with 4 SLE network hub genes are in common across naïve B cells, CD14+ and CD16+ monocytes, and pDCs. The arcs in the top half of each plot represent cells of interest, the arcs in the bottom half represent groups of co-expressed genes. The chords indicate shared co-expression groups. In each plot, the orange (denoted Bn:CD14:CD16:pDC) chords represent the genes shared across all four cell types. (**A**) Genes highly correlated with ATG5 expression. (**B**) Genes highly correlated with IFIH1 expression. (**C**) Genes highly correlated with IRAK1 expression. (**D**) Genes highly correlated with UBE2L3 expression.

## Discussion

We showed that unbiased and unsupervised training of ldVAEs on case-control scRNA-seq data robustly and consistently identifies key dysregulated processes in peripheral blood immune cells of SLE patients having a flare up. Moreover, we demonstrated that genes implicated by ldVAE case-control analysis include many differentially expressed genes and also implicate additional informative and plausible SLE-associated genes.

Across the four SLE datasets and immune populations that we analyzed, less than 20% of SLE-risk genes were differentially expressed compared to HD. Among these, IFIH1 is notable for being a correlation-network hub in healthy B cells, monocytes, and DCs that is upregulated in all these cells during SLE flares. IFIH1-mediated signaling increases IFN1 signaling and is also activated by IFN1 signaling ^40^ (Supplementary Fig. 9A). Moreover, increased expression of IFIH1 was previously shown to facilitate the onset of autoimmunity ^41^. These findings suggest that IFIH1 inhibition may act in multiple immune cell types to suppress SLE flares.

Of the three hub genes at the center of our SLE networks, ATG5 is important for autophagic vesicle formation as well as antigen presentation ^42^. The other two genes highlighted by our analysis are IRAK1 and UBE2L3. Both of these genes act downstream of TLR signaling to activate NF-κB and may contribute to upregulation of IFIH1 in SLE. IRAK1, a mediator of TLR ^43^ and IFNα/β signaling ^44^ resides on the X-chromosome ^45^, thus providing a connection to XCI escape. An SLE risk-allele of UBE2L3 increases its expression in B cells and monocytes and drives greater NF-κB mediated immune activity ^46^, and inhibition of UBE2L3 activity suppressed the response to TLR7 activation in ex-vivo human B cells ^47^. Taken together, these observations suggest that dysregulated IRAK1 and UBE2L3 activity may contribute to the emergence of a self-reinforcing signaling feedback loop thought to underlie SLE pathogenesis (Supplementary Fig. 9B).

### Study limitations

A key previous report implicated the DN2 subset of extra-follicular activated naïve B cells as key drivers of SLE pathogenesis ^18^. In SLE, pDCs up-regulate IFNα production in response to immune complexes containing IgG and nucleic acids from dying/damaged cells ^48^. Activated naïve DN2 B-cells produce the highest levels of IgG among SLE anti-dsDNA producing B cells ^49^, making them important drivers of SLE pathogenesis. These observations are in close agreement with our results showing that SLE pDCs and naïve B cells have upregulated IFNα/β responses. However, we could not reliably detect the DN2 B-cell subset in scRNA-seq data. Thus, whether naïve B cells as a whole or just the DN2 subset of naïve B cells drive SLE pathogenesis is unexplored in our analyses.

Another limitation of this study is that it is observational. Although our analysis of SLE gene co-expression network hubs implies that dysregulated debris-clearance acts upstream of IFN signaling to initiate SLE flares, perturbation experiments will be needed to establish causal relationships.

## Acknowledgements

We thank Gracie Gordon, Richard Perez, Yang Sun, and Jimmie Ye for their helpful and prompt responses to our queries about their data. We are also grateful to Peter Linsley (BRI Center for Systems Immunology) for feedback on the manuscript, and to Brian Snelgrove from the BRI High Performance Computing Team for outstanding support.

## Methods

### Data pre-processing

All single cell data were filtered with the following selection criteria:

|  |  |
| --- | --- |
| seurat_pbmc_type_score | > 0.5 & |
| nFeature_RNA | > 500 & |
| nCount_RNA | < 15E3 & |
| n_genes | < 4000 & |
| mitoRatio | < 30 & |
| meta.data\$n_reads | < 80E3 & |
| n_umis | < 15E3 |

Count data from GSE190992 and GSE214546 were normalized using scTransform ^50^. For the remaining studies, we used the normalized, batch-corrected expression values provided by the authors.

For ldVAE training and testing, the Perez et al ^27^ data was downloaded from GSE174188 as a batch-normalized Scanpy (https://scanpy.readthedocs.io) AnnotatedData (https://anndata.readthedocs.io/) object and subset to 55,120 blood immune cells from SLE donors undergoing a flare, and 486,418 blood immune cells from, healthy donors. ldVAE requires integer-valued counts as input. For input to ldVAE, we generated approximate pseudo-counts by raising the scanpy normalized expression values to the power of two, multiplying by 10, and rounding.

### Rational for use of ldVAE

The linearly decoded variational autoencoder was originally developed in the laboratories of Nir Yosef and Lior Pachter by Valentine Svensson and Adam Gayoso in 2020 ^12^. It is freely available and easy to implement via the SCVI-tools python library ^51^. However, the original short paper describing ldVAE focused on the interpretability of its latent space and its decomposition of expression data into biologically meaningful latent factors (groups of genes with correlated expression) and did not explore case-control studies.

The latent dimensions of VAEs approximate features that disentangle the underlying causes of variation in the data ^52^. The latent dimensions of ldVAE have the additional property that they each approximate a linear weighted sum of gene expression levels. We reasoned that this representation allows straight-forward interpretation of the key drivers of differences in the locations of healthy versus diseased cells. To our knowledge, to date ldVAEs have not been used to compare healthy versus disease scRNA-seq data in this way.

### ldVAE network configurations

The configurations of the 10 ldVAE models explored here are described in **Figure 4B** and Supplementary Figure 2. We found that smaller networks, especially those with fewer latent dimensions produce less granular models of the data. Notably, a single-layer encoder comprising just 64 hidden neurons and 4 latent dimensions still unambiguously captured the SLE versus HD differences. Larger networks resulted in larger numbers of cell clusters. Models larger than the ones described here tended to overfit the training set and were not explored further.

### ldVAE training and run time parameters

For all models reported, the learning rate was set to 5e-3 and the number of epochs was 250. All other training parameters (drop-out rate, etc.) were left at SCVI-tools defaults. Training curves for all 10 runs are shown in Supplementary Figure 10. Note that in all cases the validation ELBO values (normalized input-reproduction error) are lower than training ELBO values because training includes drop-out and batches whereas validation does not.

All ldVAE training was performed using a single NVIDIA GeForce RTX 2060 SUPER GPU with 8GB of dedicated GPU memory on a Windows 10 PC running WSL2 Ubuntu 22.04.3 LTS. Each training run of 250 epochs typically took approximately 2 hours to complete.

### Ad-hoc integrative network analysis

To construct a ground-truth of correlated gene expression patterns in healthy and SLE cells, we reasoned that integrating data from multiple SLE studies would better capture SLE heterogeneity. To this end, we integrated peripheral blood immune cell data from 4 SLE studies covering 237 donors, as well as 3 additional healthy-donor studies covering 1,285 donors (summarized in Supplementary Figure 1). To overcome the limited resolution of single-cell RNA-sequencing (scRNA-seq), we included data from studies using bulk RNA-seq of specific pre-sorted immune populations. To improve the sensitivity of our correlation calculations, we used the MAGIC missing-value inference method ^53^ to correct for drop out in scRNA-seq (see Supplementary Figure 11).

We used randomized label-swapping to identify statistically significant expression correlations among all gene pairs in healthy donors. We then identified gene expression modules in healthy immune cells by looking for networks of genes connected by high levels of expression correlation (Pearson’s r > 0.5). In contrast to other module detection methods such as WGCNA ^7^, our approach retains overlaps among gene expression modules, allowing us to identify functionally related modules.

To focus on SLE associated processes, we filtered our gene co-expression networks for interactions involving 126 published ^36^ GWAS risk genes and their correlations to genes reported to be differentially expressed in SLE in any of the four scRNA-seq datasets.

## Resource Availability

### Lead contacts

All scientific inquiries should be addressed to the corresponding authors Hamid Bolouri or Adam Lacy-Hulbert. Inquiries relating to the deposited data and code should be addressed to the BRI Data Manager Stephanie Osmond.

### Materials availability

No new materials were generated by this study.

### Data and code availability

This manuscript re-analyzes seven published data sets, all of which are publicly and freely available via the cited publications.

The ten ldVAE trained models described in the manuscript were saved as SCVI-Tools model objects and are available via Zenodo (DOI 10.5281/zenodo.10771142) along with an AnnotatedData object of the full training/test data, which can be read directly into the model using the provided code.

Two iPython Notebook containing all code necessary to generate the results from scratch or re-run the saved models are available via GitHub (https://github.com/hamid-bolouri/ldVAE-SLE). Additional R scripts for investigator-led network construction and integrative analysis are also available via the above GitHub repository.

