## Supplementary Figures for "Unsupervised generative AI discovers pan-leukocyte dysregulated pathways in single-cell lupus data"

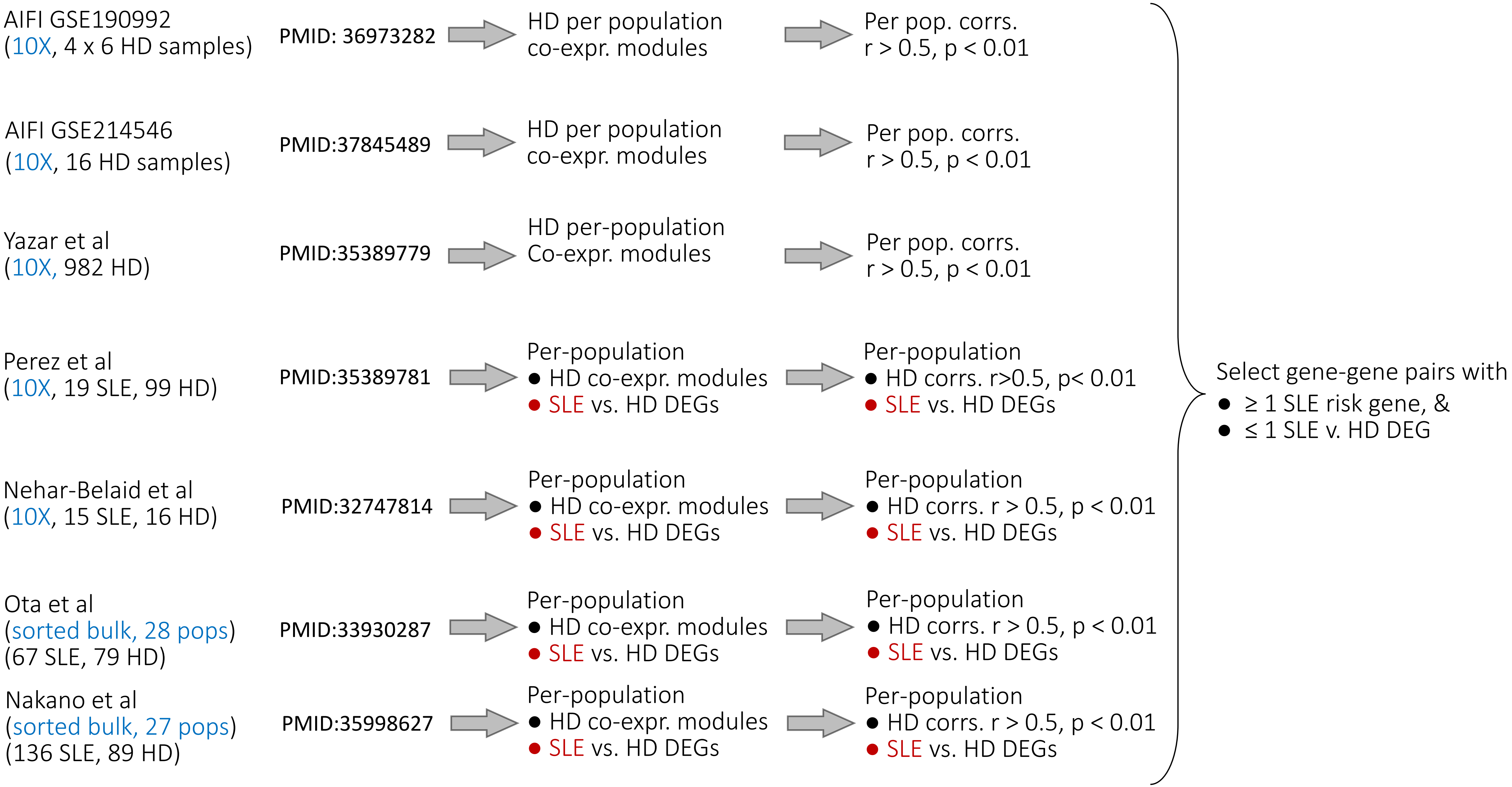
AIFI GSE190992
(10X, 4 x 6 HD samples)
PMID: 36973282
HD per population
co-expr. modules
Per pop. corrs.
r > 0.5, p < 0.01
AIFI GSE214546
(10X, 16 HD samples)
PMID:37845489
HD per population
co-expr. modules
Per pop. corrs.
r > 0.5, p < 0.01
Yazar et al
(10X, 982 HD)
PMID:35389779
HD per-population
Co-expr. modules
Per pop. corrs.
r > 0.5, p < 0.01
Perez et al
(10X, 19 SLE, 99 HD)
PMID:35389781
Per-population
● HD co-expr. modules
● SLE vs. HD DEGs
Per-population
● HD corrs. r>0.5, p< 0.01
● SLE vs. HD DEGs
Nehar-Belaid et al
(10X, 15 SLE, 16 HD)
PMID:32747814
Per-population
● HD co-expr. modules
● SLE vs. HD DEGs
Per-population
● HD corrs. r > 0.5, p < 0.01
● SLE vs. HD DEGs
Ota et al
(sorted bulk, 28 pops)
(67 SLE, 79 HD)
PMID:33930287
Per-population
● HD co-expr. modules
● SLE vs. HD DEGs
Per-population
● HD corrs. r > 0.5, p < 0.01
● SLE vs. HD DEGs
Nakano et al
(sorted bulk, 27 pops)
(136 SLE, 89 HD)
PMID:35998627
Per-population
● HD co-expr. modules
● SLE vs. HD DEGs
Per-population
● HD corrs. r > 0.5, p < 0.01
● SLE vs. HD DEGs
Select gene-gene pairs with
● ≥ 1 SLE risk gene, &
● ≤ 1 SLE v. HD DEG

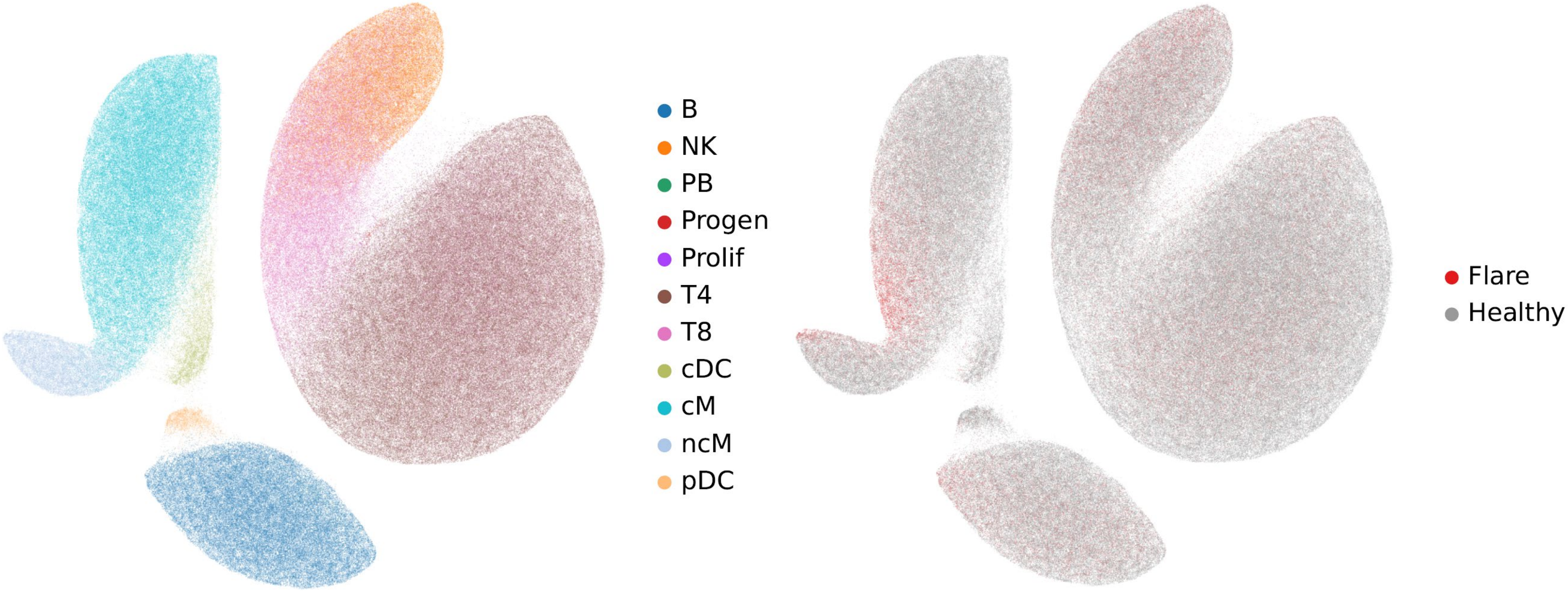


Supplementary Fig. 2A.

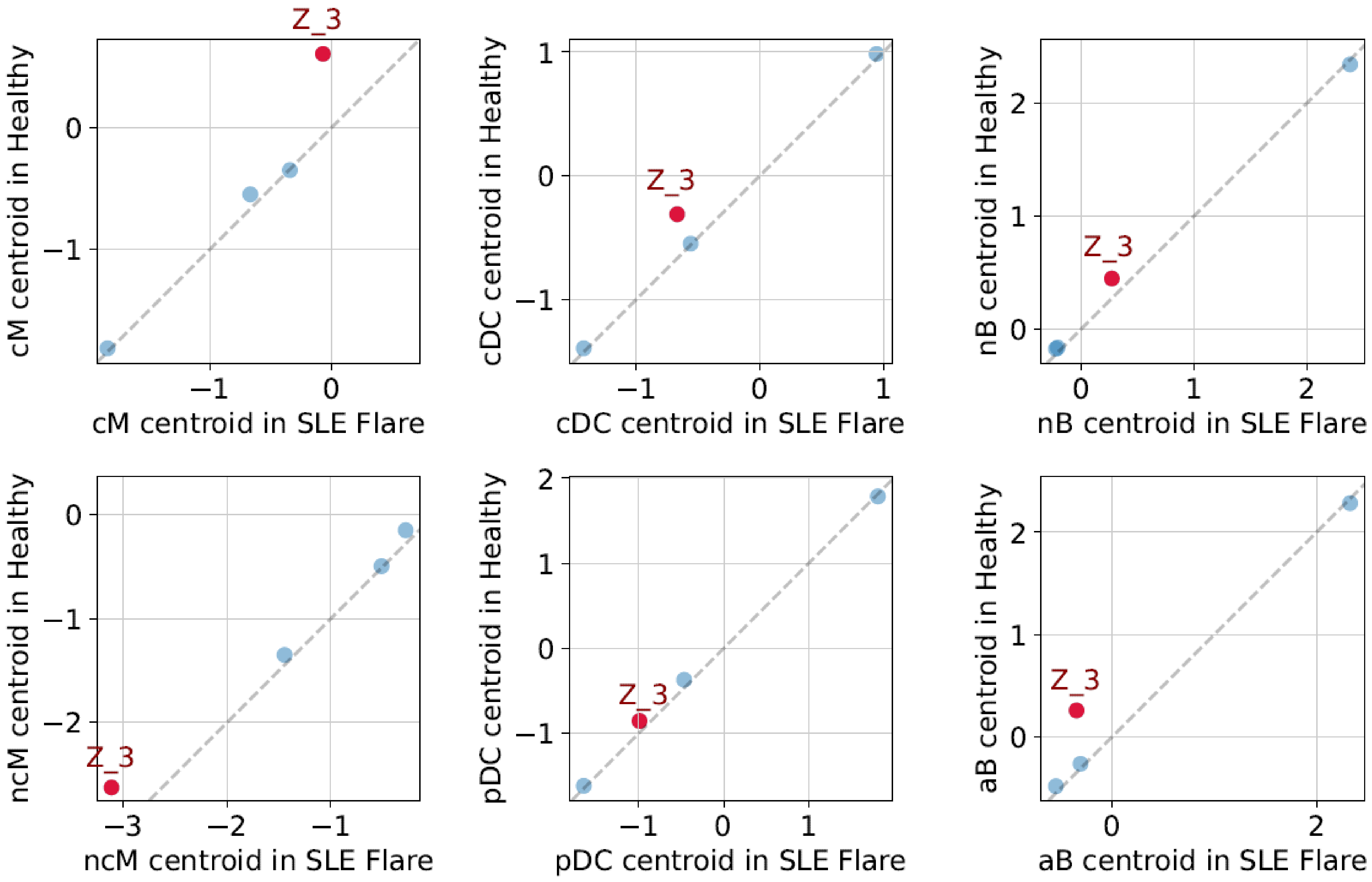


Supplementary Fig. 2B.

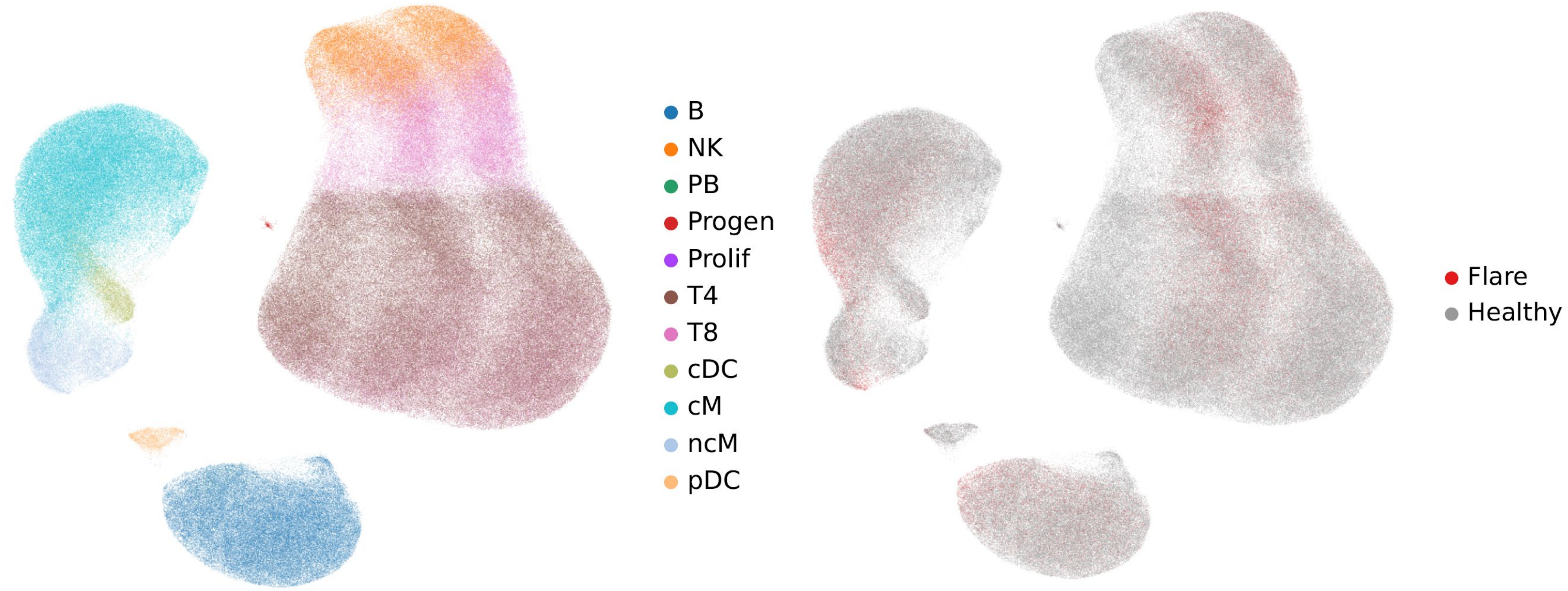


Supplementary Fig. 2C.

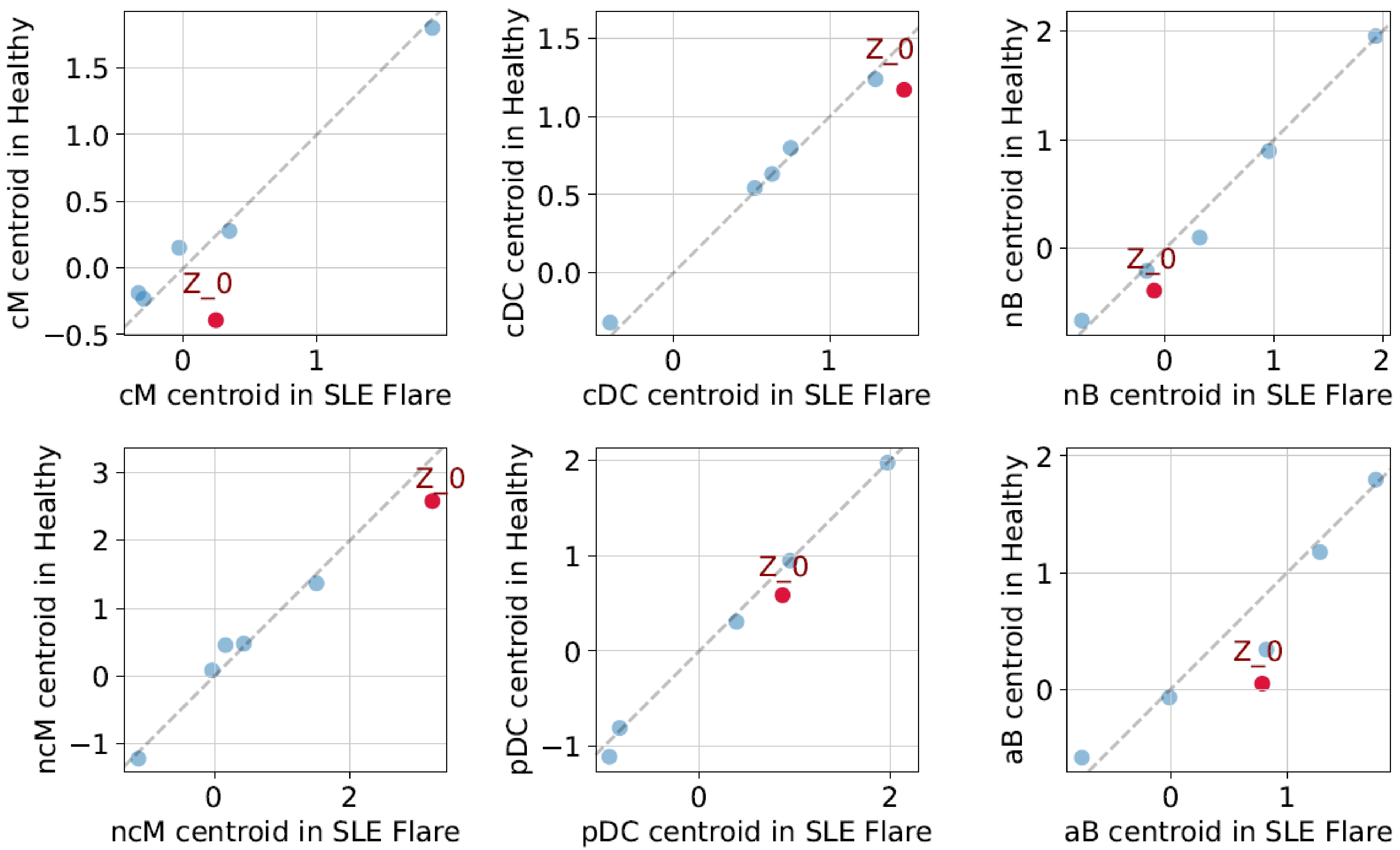


Supplementary Fig. 2D.

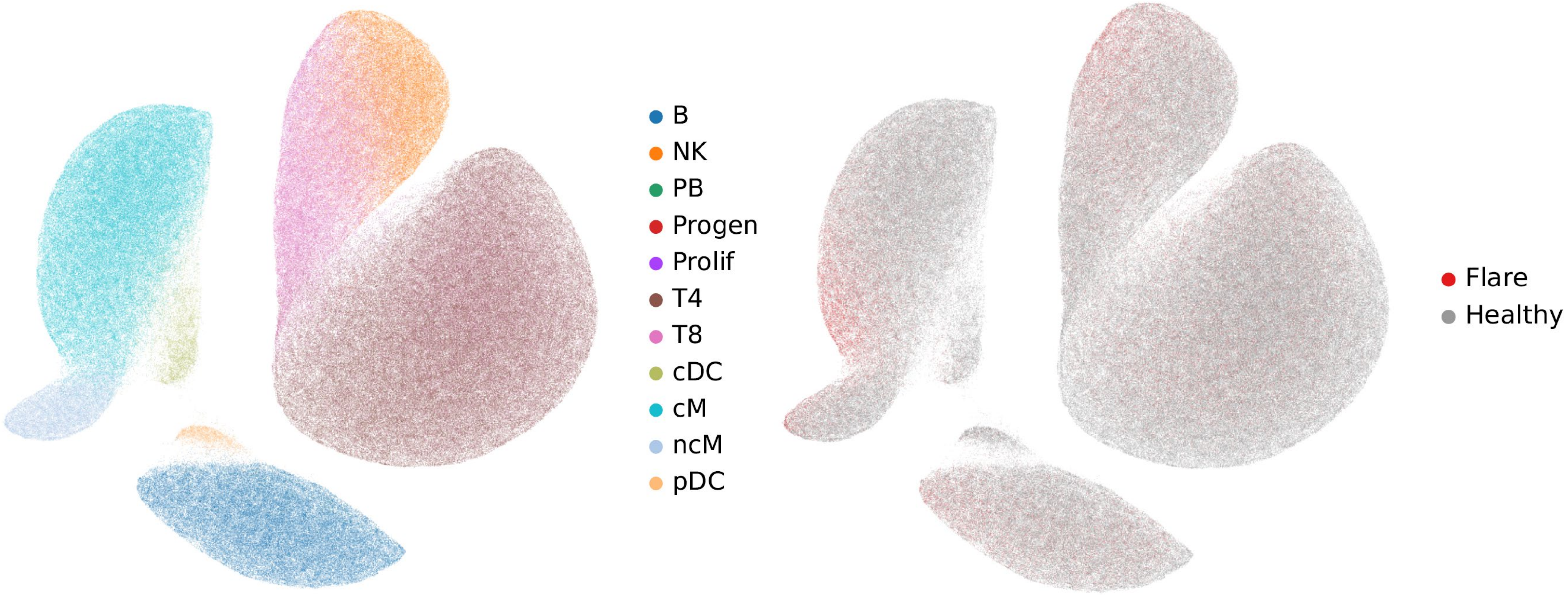


Supplementary Fig. 2E.

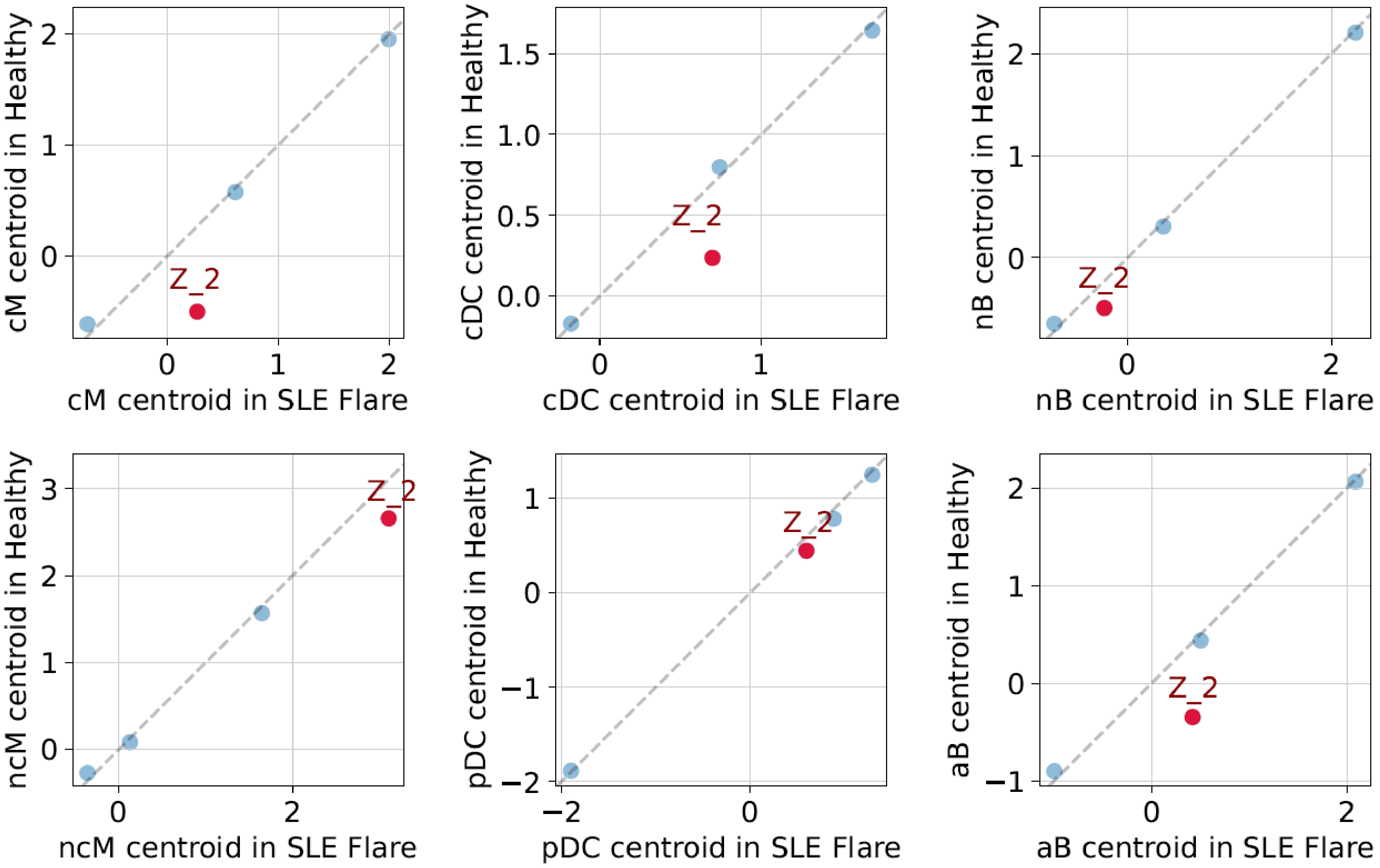


Supplementary Fig. 2F.

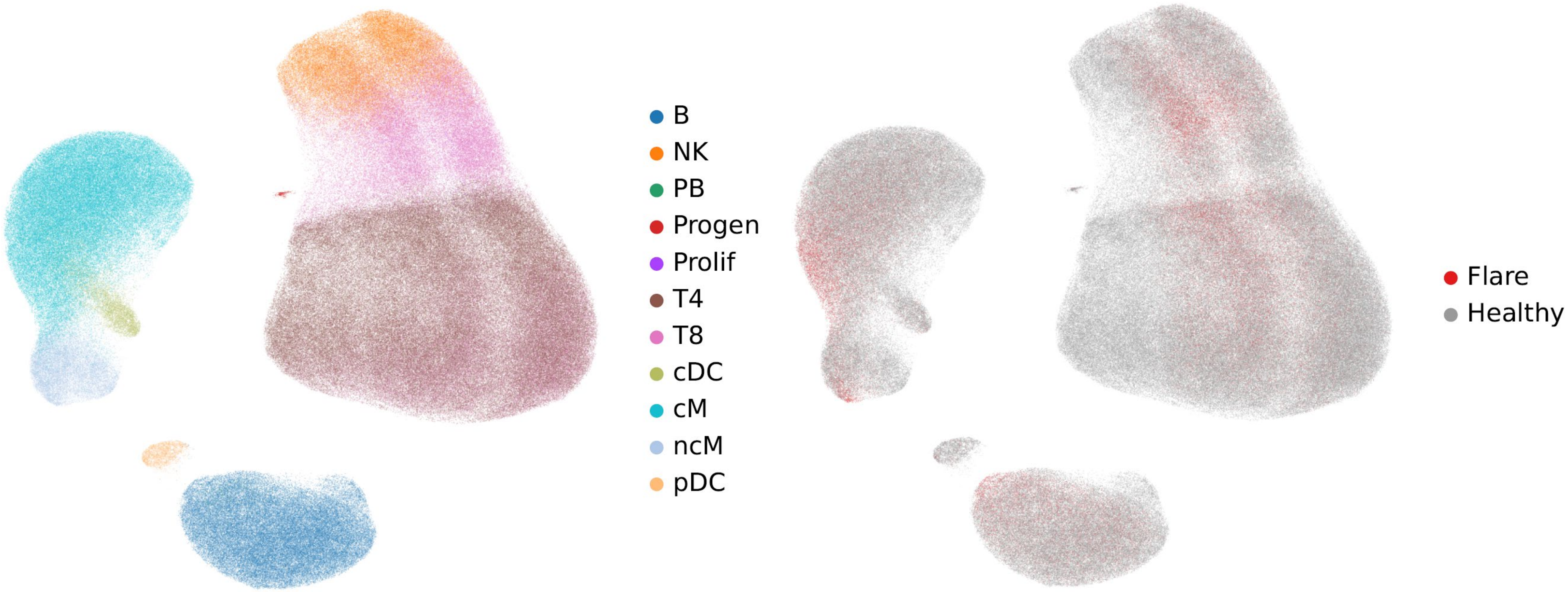


Supplementary Fig. 2G.

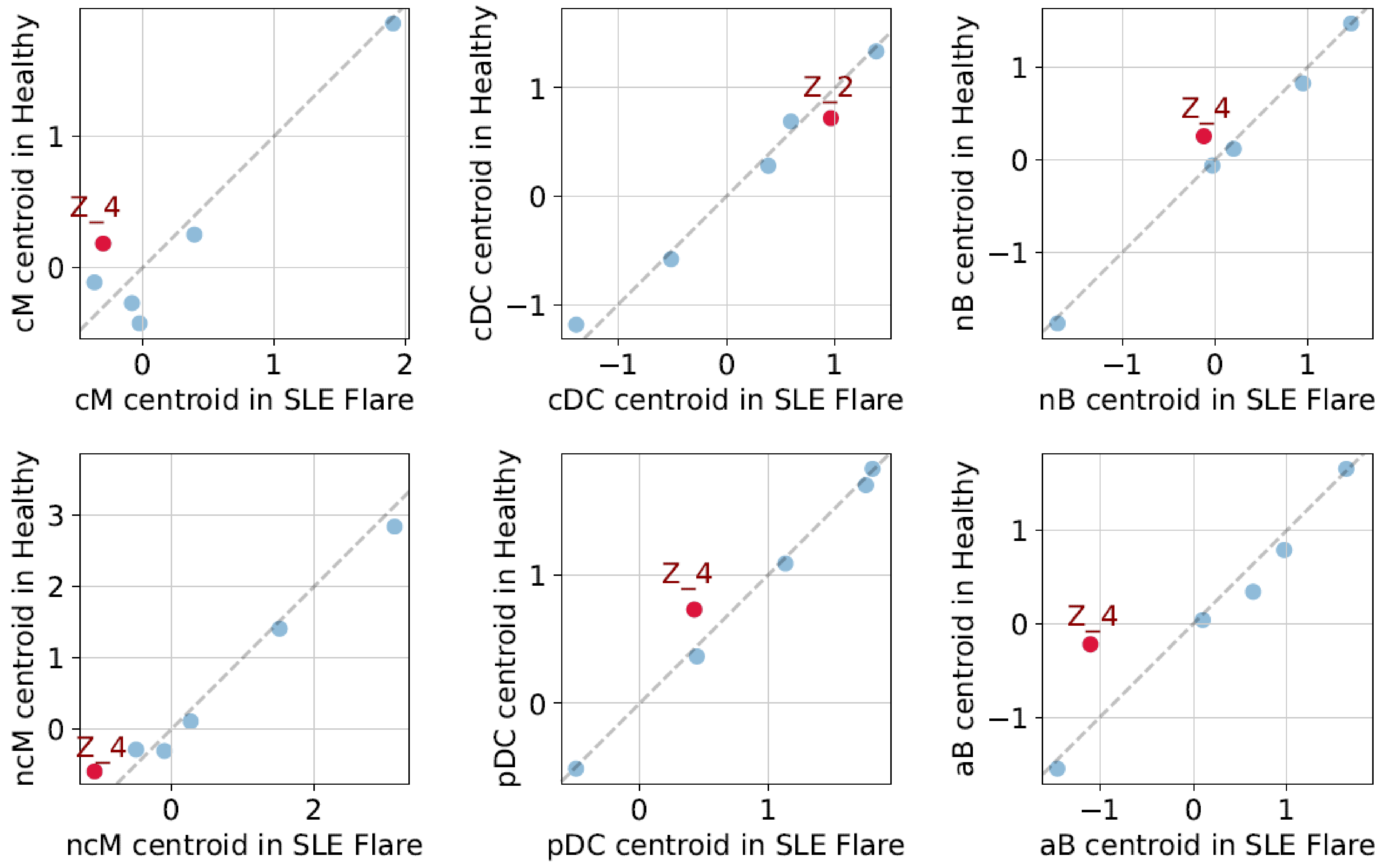


Supplementary Fig. 2H.

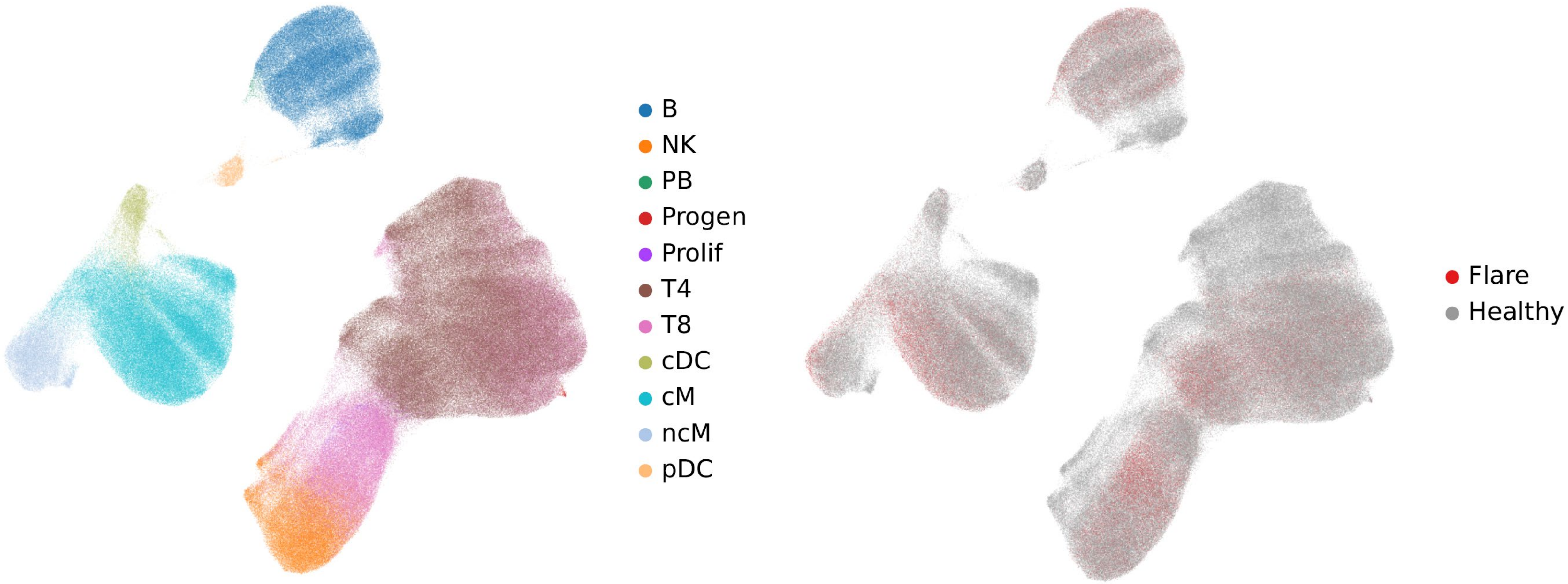


Supplementary Fig. 2I.

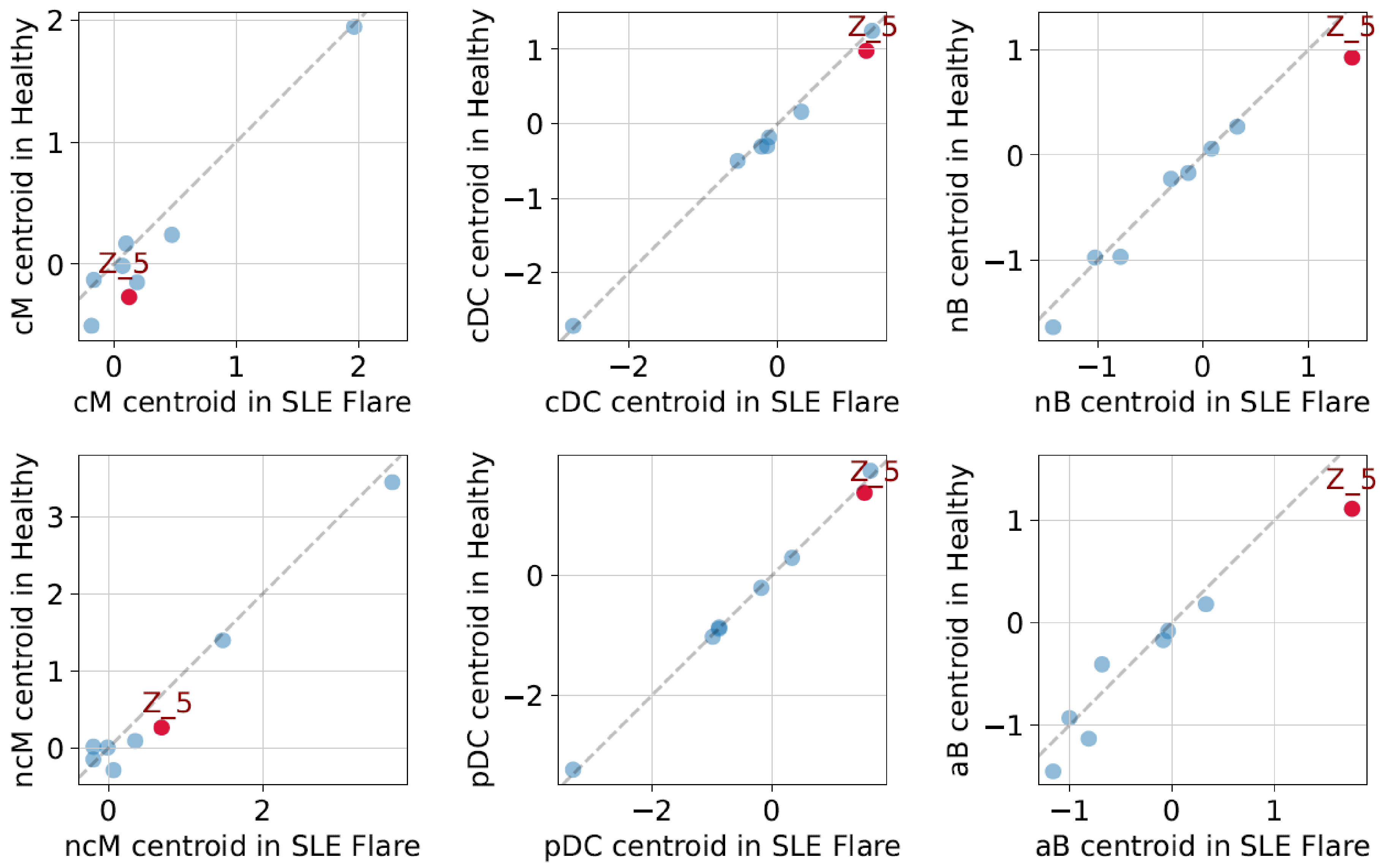


Supplementary Fig. 2J.

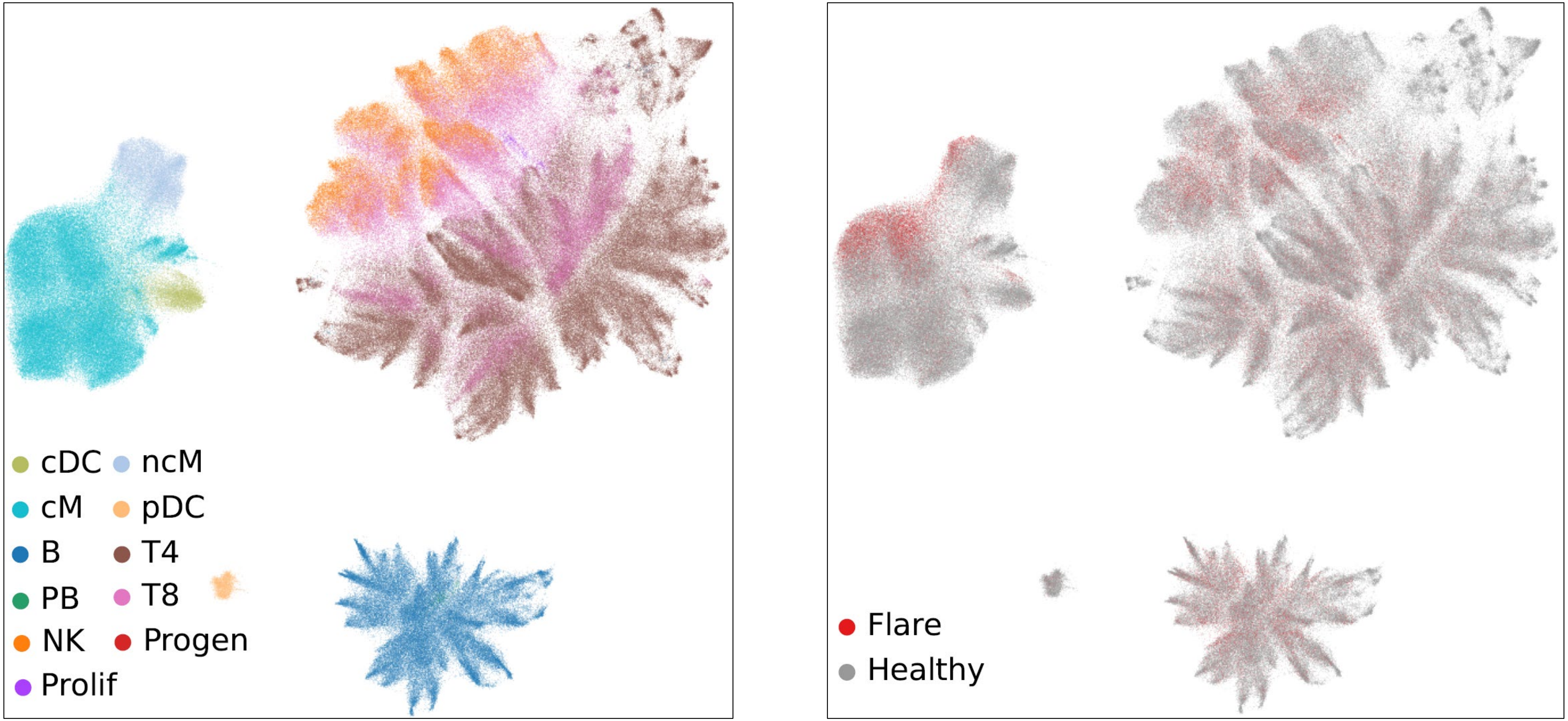


Supplementary Fig. 2K.

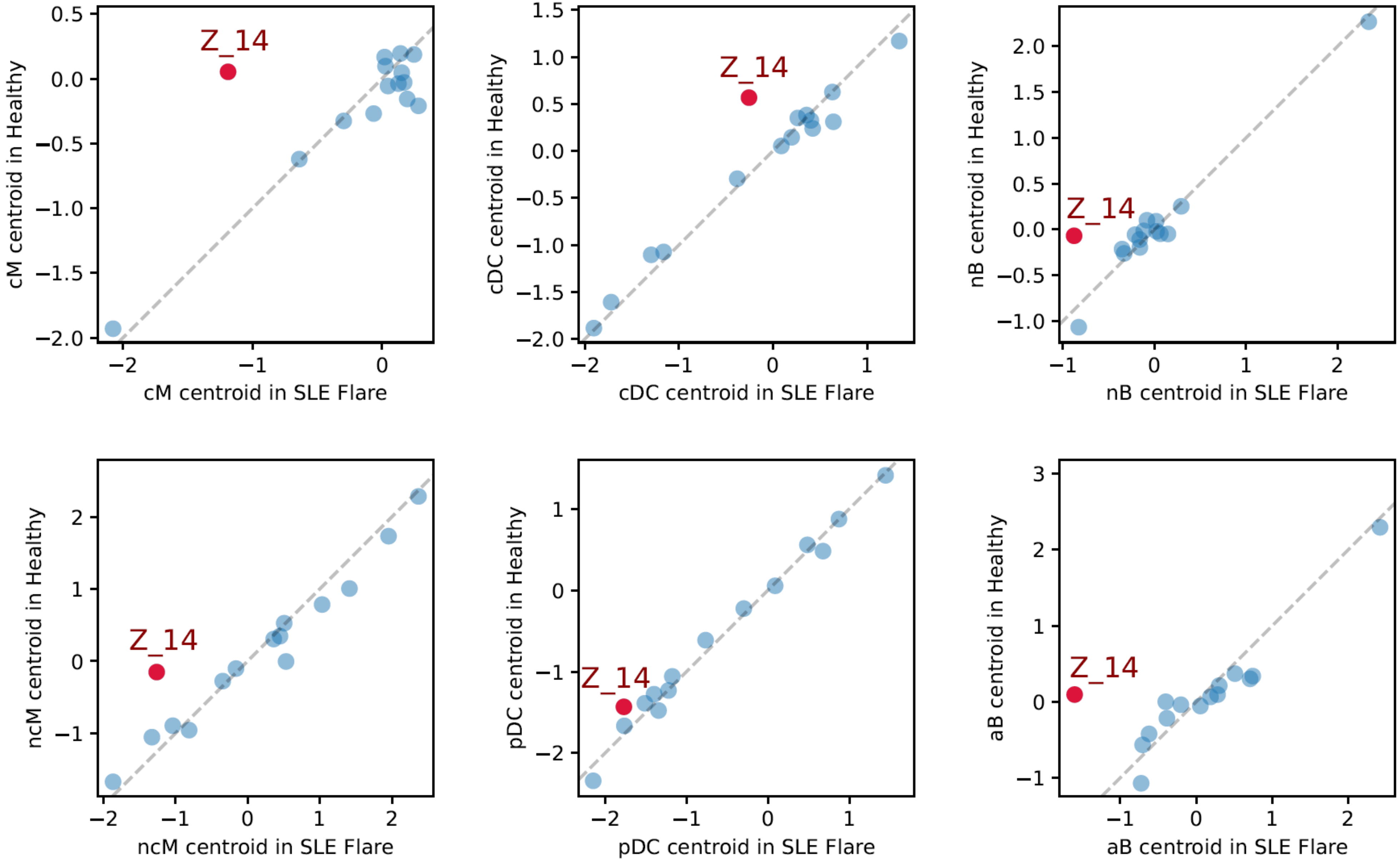


Supplementary Fig. 2L.

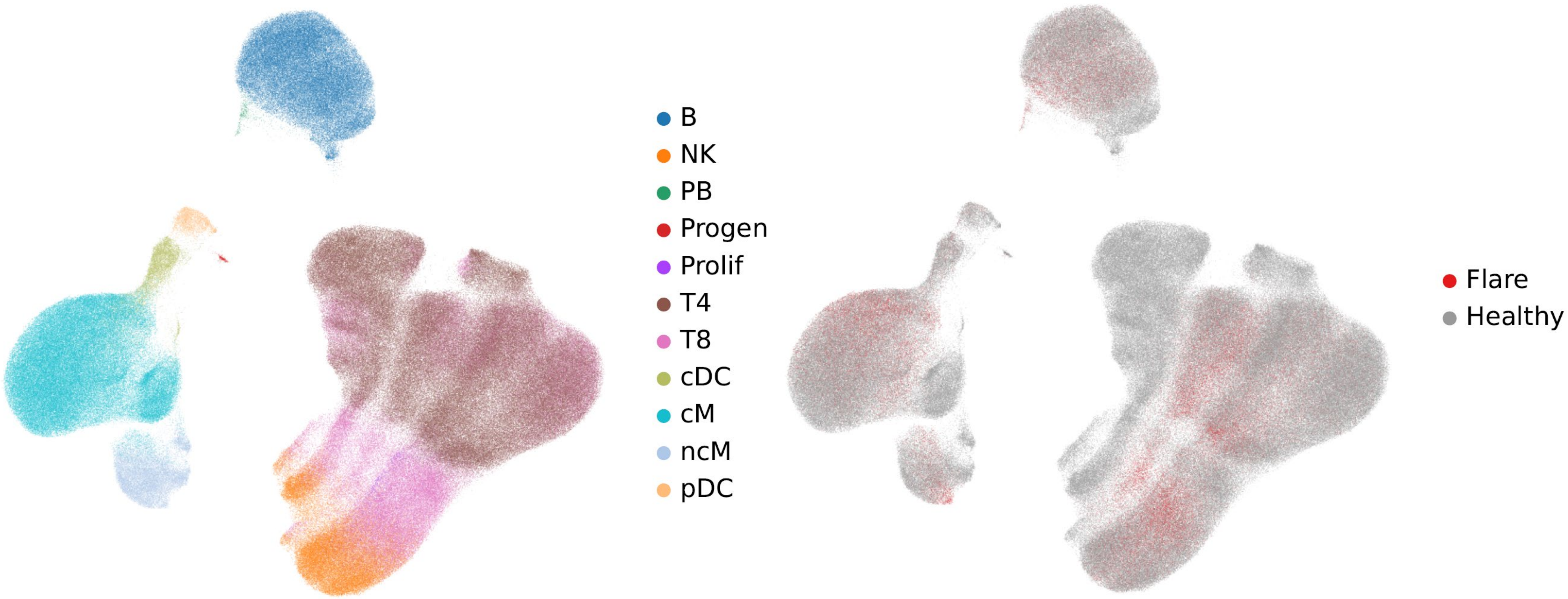


Supplementary Fig. 2M.

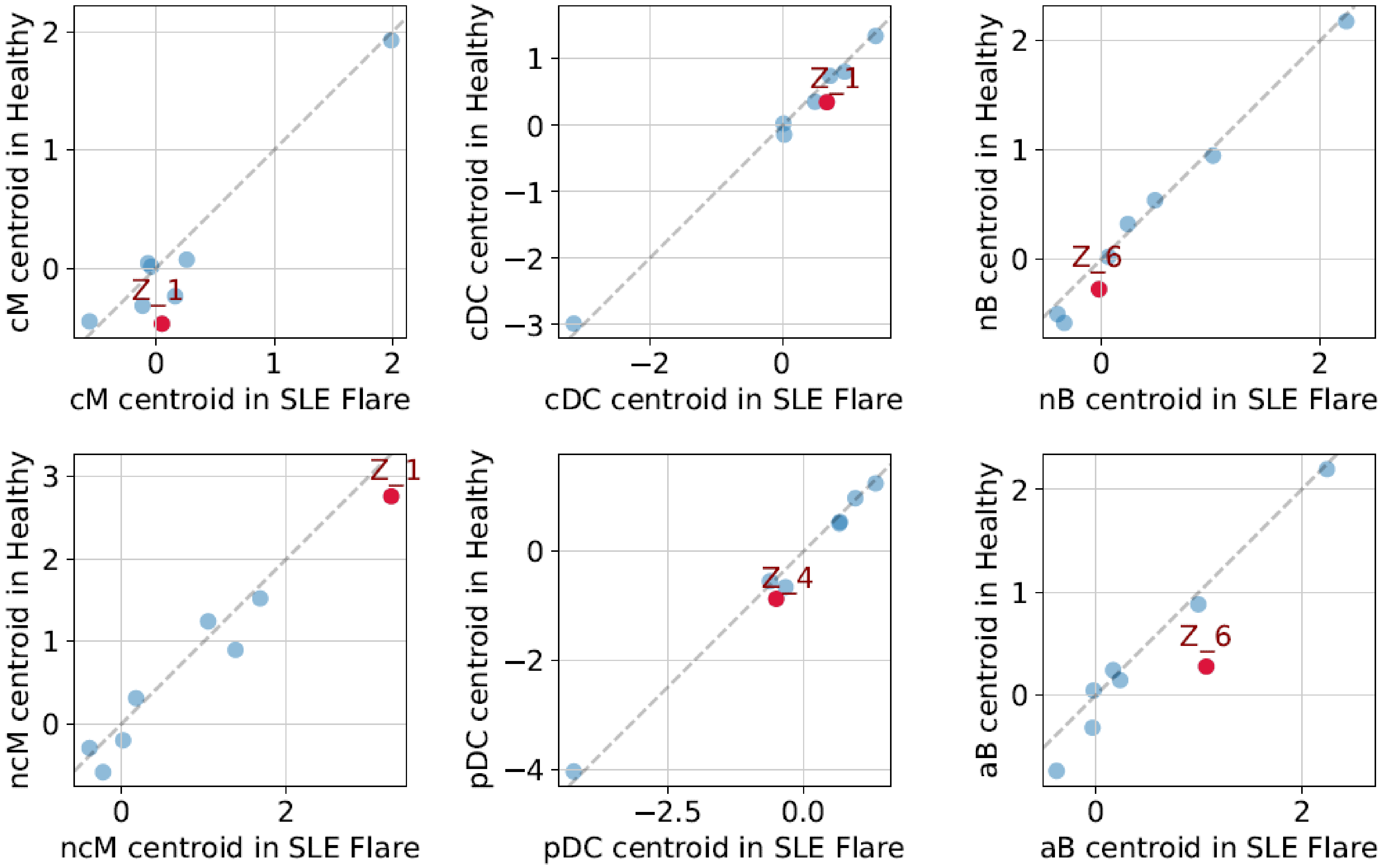


Supplementary Fig. 2N.

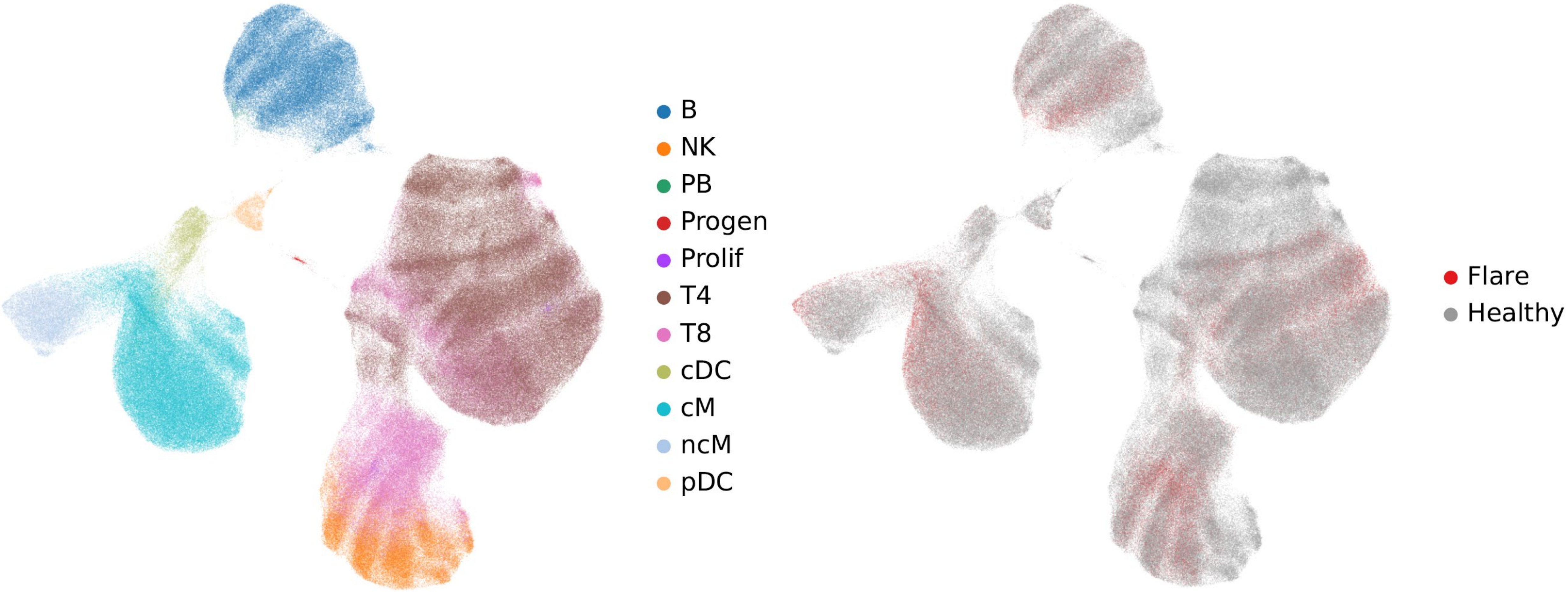


Supplementary Fig. 2O.

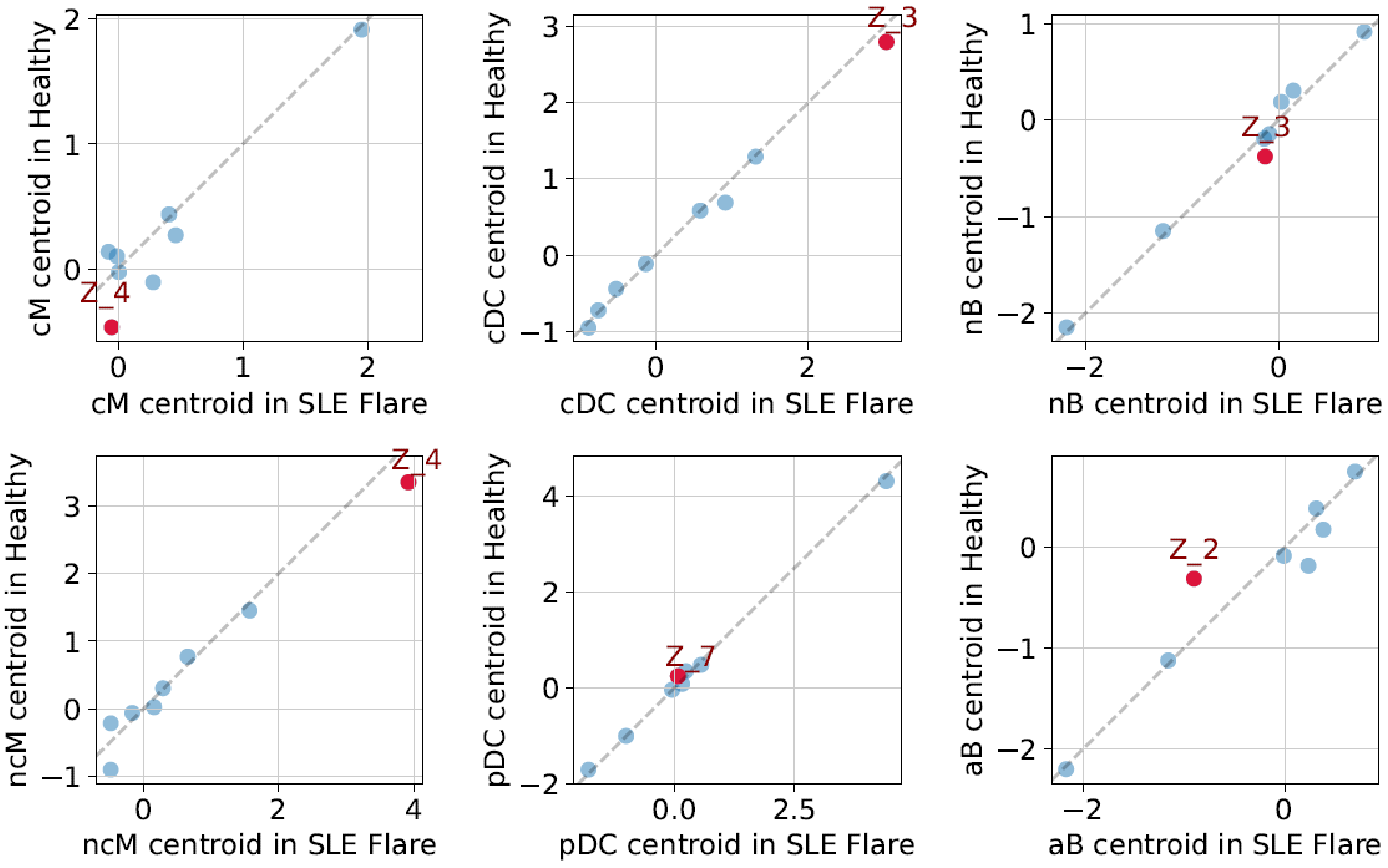


Supplementary Fig. 2P.

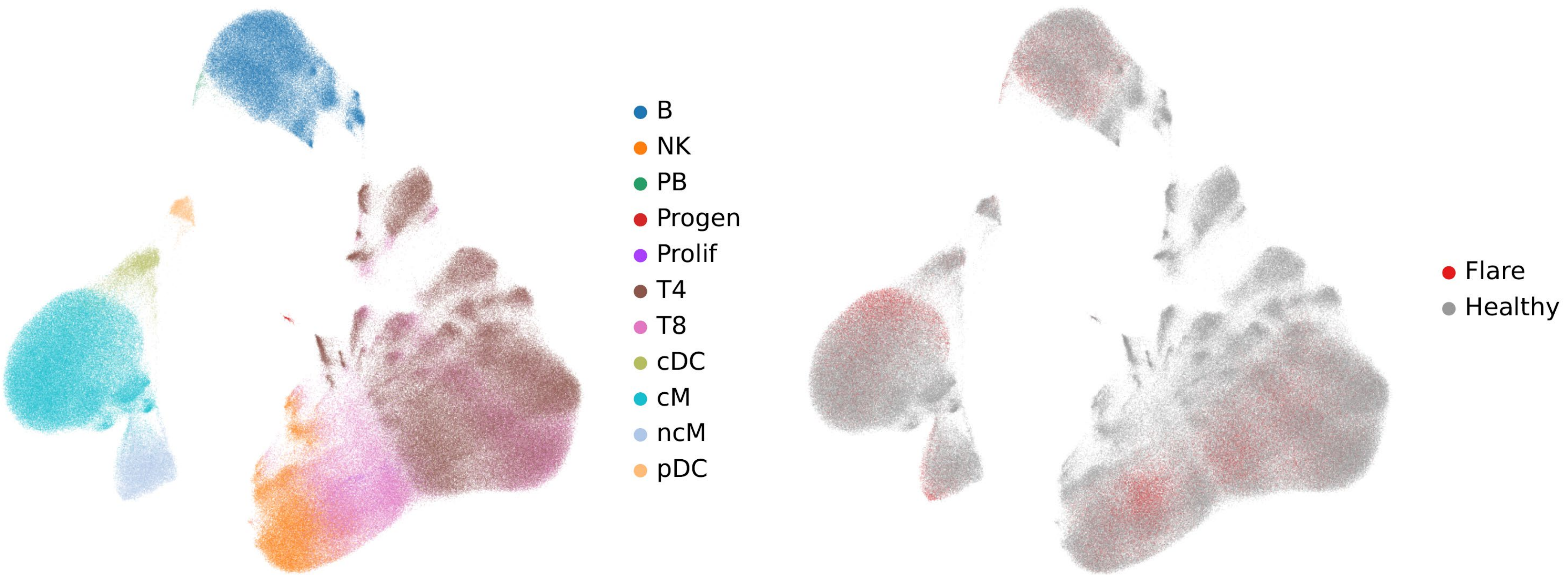


Supplementary Fig. 2Q.

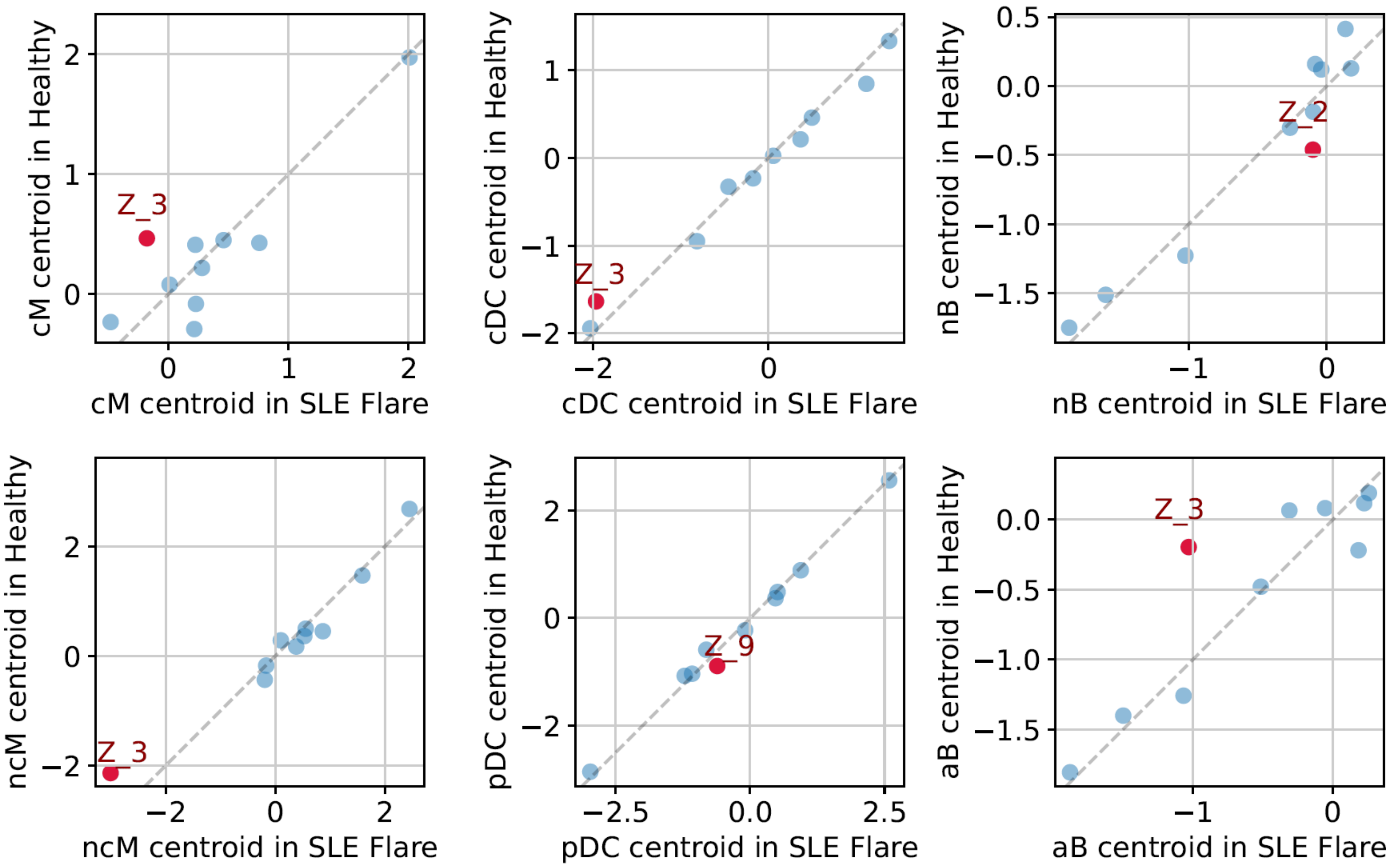


Supplementary Fig. 2R.

#### Model L1H128Z15

| Top 25 genes with higher expression along Z14 in SLE | | |
|---|---|---|
| **Gene** | **Weight** | **Notes** |
| **EGR1** | -0.21 | Blunts the inflammatory response PMC7806227 |
| **HERC5** | -0.37 | IFN signaling |
| **HSH2D** | -0.24 | SLE IFN signature gene, PMC5937945 |
| **IFI27** | -1.74 | IFN signaling |
| **IFI44** | -2.06 | IFN signaling |
| **IFI44L** | -1.67 | IFN signaling |
| **IFI6** | -0.99 | IFN signaling |
| **IFIT1** | -0.36 | IFN signaling |
| **IFIT2** | -0.27 | IFN signaling |
| **IFIT3** | -2.48 | IFN signaling |
| **IFITM3** | -0.28 | IFN signaling |
| **IRF7** | -2.08 | IFN signaling |
| **ISG15** | -0.87 | IFN signaling |
| **LY6E** | -0.22 | ISG. Limits mono activation PMID: 25225669 |
| **ITGAM** | -0.22 | LC3-associated phagocytosis, PMC6103671 |
| **MT2A** | -0.22 | Suppresses apoptosis PMC5465913 |
| **NCF1** | -0.24 | Regulates IFN1 & phagosome proteolysis PMID: 34556485 |
| **PLSCR1** | -0.34 | Modulates phagocytosis, PMC4712888 |
| **PTGDS** | -0.20 | Inhibits apoptosis PMC10777022 |
| **SAMD9** | -0.21 | Is induced by IRF1, mediates apoptosis PMC3169306 |
| **SCO2** | -0.24 | Iduces apoptosis PMC3624269 |
| **CTNNB1** | -0.19 | Mediates mucosal tolerance, PMC5081350 |
| **STAT1** | -1.66 | Autoimmunity, inflammation & apoptosis PMC4920021 |
| **TCF4** | -0.20 | Suppresses autoimmune neuroinflammation, PMC8415100 |
| **THRB** | -0.33 | |

#### Model L1H128Z8

| Top 25 genes with higher expression along Z2 in SLE | | |
|---|---|---|
| **Gene** | **weight** | **Notes** |
| **CES1** | 1.67 | Up-regulated by NF-κB in inflamation, PMID: 35965153 |
| **CX3CR1** | 1.85 | Inflammatory signaling |
| **GBP5** | 2.10 | IFN signaling |
| **HERC5** | 2.23 | IFN signaling |
| **HSH2D** | 1.09 | SLE IFN signature gene, PMC5937945 |
| **IFI44** | 0.97 | IFN signaling |
| **IFIT1** | 2.52 | IFN signaling |
| **IFIT2** | 2.87 | IFN signaling |
| **IFIT3** | 2.31 | IFN signaling |
| **MXD1** | 0.94 | CD16+ mono SLE IFN sig. gene, PMC3877094 |
| **SMC4** | 1.70 | Promotes innate inflammatory response, PMID: 29803706 |
| **CCDC50** | 1.20 | Autophagy receptor |
| **CD163** | 0.92 | endocytosis |
| **QPCT** | 0.97 | Inhibits phagocytosis, PMC8932921 |
| **S1PR5** | 1.11 | Higher expression reduces efferocytosis, PMID: 27868302 |
| **STOM** | 1.26 | Mac phagocytotsis: PMC9942578 |
| **ZBP1** | 2.28 | binds DNA, induces IFN1 & PANoptosis, PMC9499459 |
| **AL928742.12** | 1.03 | Down-regulated in MS, PMID: 31576494 |
| **APOL6** | 0.89 | UP-regulated in RA, PMID: 28882568 |
| **CEP78** | 1.72 | Marker of immune tolerance, PMC4361657 |
| **GPRIN3** | 2.71 | May be RA ssociated PMC3396452 |
| **IL2RB** | 1.52 | Cellular immune response |
| **TGFBR3** | 2.08 | Marker of membranous lupus nephritis, PMC8676385. Marker of immune tolerance, PMC4361657 |
| **C1orf21** | 0.95 | |
| **MEGF9** | 0.96 | |

#### Model L1H128Z6

| Top 25 genes with higher expression along Z4 in SLE | | |
|---|---|---|
| **Gene** | **weight** | **Notes** |
| **CD84** | -1.33 | IFN signaling |
| **GBP5** | -1.31 | IFN signaling |
| **HERC5** | -1.55 | IFN signaling |
| **HSPA1A** | -1.67 | IFN signaling |
| **HSPA1B** | -0.67 | IFN signaling |
| **IFI27** | -0.88 | IFN signaling |
| **IFI44** | -0.80 | IFN signaling |
| **IFIT1** | -1.50 | IFN signaling |
| **IFIT3** | -1.18 | IFN signaling |
| **PTMS** | -0.89 | Modulates PMC397191 pro-IFN1 effects of ProTα PMC7126549 |
| **ATG2A** | -0.90 | Autophagy related 2A |
| **CDKN1A** | -0.60 | Autophagy PMC6770903 |
| **FAM129A** | -1.52 | Regulates autophagy & apoptosis, PMC9062034 |
| **FAM46C** | -0.93 | Cytotoxic. Drives IRF4 PMC5586597 |
| **GZMK** | -0.61 | Cytotoxicity, infalmmation PMC8129556 |
| **MIR4435-1HG** | -0.69 | Regulates apoptosis PMC9205143 |
| **SAP30** | -1.28 | Regulates apoptosis PMC10781503 |
| **STOM** | -0.87 | Mac phagocytotsis: PMC9942578 |
| **TGFBR3** | -1.07 | Marker of membranous lupus nephritis, PMC8676385 & immune tolerance, PMC4361657 |
| **BHLHE40** | -0.78 | Multiple autoimmune roles PMC7606821 |
| **CES1** | -0.59 | Regulates macrophage IL1β PMID: 3734804 |
| **LMNA** | -0.66 | Myeloid differentiation, activation PMC7504305 |
| **MYBL1** | -0.87 | B-cell survival PMID: 10910917 |
| **PRDM1** | -0.94 | Self-ractive plasma cells PMC6331720 |
| **RGS1** | -0.59 | Role in CD8+ TRM activation PMC10158727 |

| Top 25 genes with higher expression along Z2 in SLE | | |
|---|---|---|
| **Gene** | **weight** | **Notes** |
| **APOBEC3A** | 0.39 | IFNα response PMC2118075, PMC10457583. Autoimmune PMID:38060252. DNA damage PMC3090015 |
| **CX3CR1** | 0.91 | IFN signaling |
| **HSH2D** | 0.36 | IFN signaling |
| **IFIT2** | 0.55 | IFN signaling |
| **IFIT3** | 0.57 | IFN signaling |
| **IRF7** | 0.39 | IFN signaling |
| **RHOC** | 0.66 | IFNγ responsive PMC6686045 |
| **C1QA** | 0.46 | efferocytosis PMC5173410, lupus PMC8082710 |
| **C1QB** | 0.37 | efferocytosis, lupus |
| **CDKN1C** | 0.41 | Has both pro- and anti-apoptotic functions PMC7575724 |
| **IGFBP7** | 0.53 | Mediates senescence PMC9773704 |
| **ASCL2** | 0.59 | Enhances antibody production PMID: 3407772 |
| **CEP78** | 0.49 | Marker of immune tolerance, PMC4361657 |
| **FCGR3A** | 0.81 | Lupus associated |
| **FCGR3B** | 0.34 | Lupus associated |
| **HMOX1** | 0.56 | Has immuno-suppressive effects PMC7308058 |
| **MEG3** | 0.38 | Promotes inflammation PMID33189612. Suppresses Tregs PMC8212454 |
| **MS4A7** | 0.41 | Short isoform regulates mac polarization PMID: 36944954 |
| **FGD2** | 0.36 | |
| **HES4** | 1.36 | |
| **LRRC26** | 0.44 | |
| **LYPD2** | 0.57 | |
| **RP11-290F20.3** | 0.40 | |
| **SCT** | 0.66 | |
| **VMO1** | 0.40 | |

Supplementary Fig. 3A.

#### Model L1H128Z4

**Top 25 genes with higher expression along Z1 in SLE**

| Gene | Weight | Notes |
|---|---|---|
| **APOBEC3A** | 0.67 | IFNa response PMC2118075, PMC10457583. Autoimmune PMID:38060252. DNA damage PMC3090015 |
| **HERC5** | 1.45 | IFN signaling |
| **IFI27** | 0.92 | IFN signaling |
| **IFI44** | 0.85 | IFN signaling |
| **IFI44L** | 0.64 | IFN signaling |
| **IFIT1** | 0.93 | IFN signaling |
| **IFIT2** | 1.47 | IFN signaling |
| **IFIT3** | 1.54 | IFN signaling |
| **RHOC** | 0.77 | IFNg responsive PMC6686045 |
| **C1QA** | 0.57 | efferocytosis, lupus |
| **CDKN1C** | 0.58 | Has both pro- and anti-apoptotic functions PMC7575724 |
| **CX3CR1** | 1.03 | efferocytosis, lupus |
| **FCAR** | 0.61 | FcαRI, phagocytosis, lupus associated |
| **TLR2** | 0.75 | Phagocytosis PMC3342043 |
| **ASCL2** | 0.66 | Enhances antibody production PMID: 3407772 |
| **CEP78** | 0.81 | Marker of immune tolerance, PMC4361657 |
| **FCGR3A** | 0.85 | lupus |
| **HMOX1** | 0.95 | Has immuno-suppressive effects PMC7308058 |
| **MS4A7** | 0.56 | Short isoform regulates mac polarization PMID: 36944954 |
| **HES4** | 1.66 | |
| **LYPD2** | 0.68 | |
| **VMO1** | 0.57 | |
| **CCR1** | 0.77 | |
| **FAM198B** | 0.79 | |
| **TRIB1** | 0.62 | |

#### Model L2H64Z8

**Top 25 genes with higher expression along Z5 in SLE**

| Gene | Weight | Notes |
|---|---|---|
| **BHLHE40** | 1.13 | up-regulates IFNγ, PMC7606821 |
| **CD84** | 1.45 | Drives IFNγ, PMID11564780. In SLE causes B & T cell hyperstimulation, PMID22549634 |
| **CX3CR1** | 1.21 | Inflammatory signaling |
| **GBP5** | 1.40 | IFN signaling |
| **HERC5** | 1.42 | IFN signaling |
| **HSH2D** | 0.72 | SLE IFN signature gene, PMC5937945 |
| **IFIT1** | 1.57 | IFN signaling |
| **IFIT2** | 2.17 | IFN signaling |
| **IFIT3** | 1.50 | IFN signaling |
| **PTMS** | 2.32 | Modulates PMC397191 pro-IFN1 effects of ProTα PMC7126549 |
| **SMC4** | 1.13 | Promotes IFNβ response, PMID: 29803706 |
| **ANKRD28** | 0.93 | PP6 subunit, regulates autophagy, PMID: 30036567 |
| **CCDC50** | 1.62 | Autophagy receptor: PMC7852694 |
| **FAM129A** | 1.97 | Regulates autophagy & apoptosis, PMC9062034 |
| **S1PR5** | 0.75 | Regulates efferocytosis, PMID: 27868302 |
| **STOM** | 0.91 | Macrophage phagocytotsis: PMC9942578 |
| **AL928742.12** | 0.70 | Down-regulated in MS, PMID: 31576494 |
| **CEP78** | 1.18 | Marker of immune tolerance, PMC4361657 |
| **HSPA1A** | 2.21 | Tolerogenic PMC3343630 |
| **HSPA1B** | 1.62 | Tolerogenic PMC3343630 |
| **NRGN** | 0.85 | Drives B & T cell tolerance, PMC9649591 |
| **TGFBR3** | 1.91 | Marker of membranous lupus nephritis, PMC8676385 & immune tolerance, PMC4361657 |
| **PTPN12** | 0.62 | Secondary T cell activation PMID: 20727793 |
| **APOBR** | 2.04 | |
| **C1orf21** | 0.80 | |

#### Model L1H64Z6

**Top 25 genes with higher expression along Z0 in SLE**

| Gene | Weight | Notes |
|---|---|---|
| **APOBEC3A** | 0.64 | IFNα response PMC2118075, PMC10457583. Autoimmune PMID:38060252. DNA damage PMC3090015 |
| **CD84** | 0.75 | Drives IFNγ, PMID11564780. In SLE causes B & T cell hyperstimulation, PMID22549634 |
| **CX3CR1** | 0.64 | Inflammatory signaling |
| **GBP5** | 0.96 | IFN signaling |
| **HERC5** | 1.20 | IFN signaling |
| **HSPA1B** | 0.76 | IFN signaling |
| **IFI27** | 0.75 | IFN signaling |
| **IFI44** | 0.65 | IFN signaling |
| **IFIT1** | 0.67 | IFN signaling |
| **IFIT2** | 1.38 | IFN signaling |
| **IFIT3** | 1.48 | IFN signaling |
| **PTMS** | 0.82 | Modulates PMC397191 pro-IFN1 effects of ProTα PMC7126549 |
| **RHOC** | 0.63 | IFNγ responsive PMC6686045 |
| **C1QA** | 0.60 | efferocytosis PMC5173410, lupus PMC8082710 |
| **FAM129A** | 0.76 | Regulates autophagy & apoptosis, PMC9062034 |
| **FCGR3A** | 0.64 | |
| **HMOX1** | 0.96 | Has immuno-suppressive effects PMC7308058 |
| **HSPA1A** | 1.02 | Tolerogenic PMC3343630 |
| **MS4A7** | 0.58 | Short isoform regulates mac polarization PMID: 36944954 |
| **TGFBR3** | 0.70 | Marker of membranous lupus nephritis, PMC8676385 & immune tolerance, PMC4361657 |
| **GPRIN3** | 1.12 | RA association PMC3396452 |
| **HES4** | 1.26 | |
| **IDO1** | 0.67 | |
| **LPAR6** | 0.58 | |
| **LYPD2** | 0.64 | |

#### Model L1H64Z4

**Top 25 genes with increased expression along Z3 in SLE**

| Gene | weight | Notes |
|---|---|---|
| **APOBEC3A** | -0.59 | IFNα response PMC2118075, PMC10457583. Autoimmune PMID:38060252. DNA damage PMC3090015 |
| **CX3CR1** | -0.88 | Interferon signaling |
| **IFI27** | -0.83 | |
| **IFI44** | -0.54 | Interferon signaling |
| **IFI44L** | -0.51 | Interferon signaling |
| **IFIT1** | -0.61 | Interferon signaling |
| **IFIT2** | -1.11 | Interferon signaling |
| **IFIT3** | -1.32 | Interferon signaling |
| **RHOC** | -0.71 | IFNγ responsive PMC6686045 |
| **C1QA** | -0.57 | efferocytosis, lupus |
| **CDKN1C** | -0.56 | Has both pro- and anti-apoptotic functions PMC7575724 |
| **FCAR** | -0.54 | FcαRI, phagocytosis, lupus associated |
| **IGFBP7** | -0.58 | Mediates senescence PMC9773704 |
| **TLR2** | -0.70 | |
| **ASCL2** | -0.57 | |
| **CCR1** | -0.67 | Autoimmunity PMID: 15202722 |
| **CEP78** | -0.57 | |
| **FCGR3A** | -0.86 | Lupus associated |
| **HMOX1** | -0.85 | |
| **MS4A7** | -0.55 | |
| **HES4** | -1.39 | |
| **LYPD2** | -0.61 | |
| **RP11-290F20.3** | -0.56 | |
| **SCT** | -0.53 | |
| **VMO1** | -0.50 | |

Supplementary Fig. 3B.

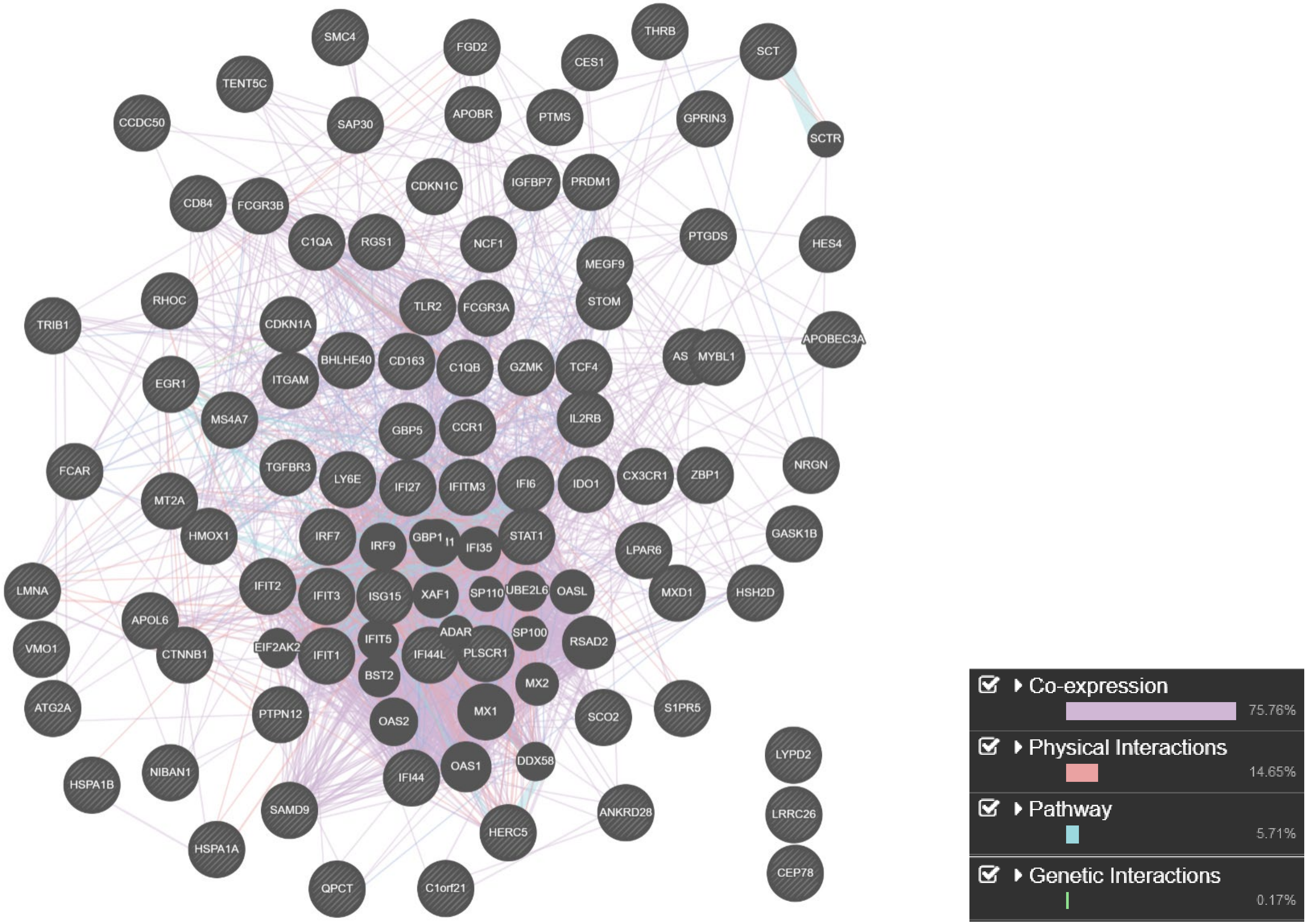


Supplementary Fig. 4A.

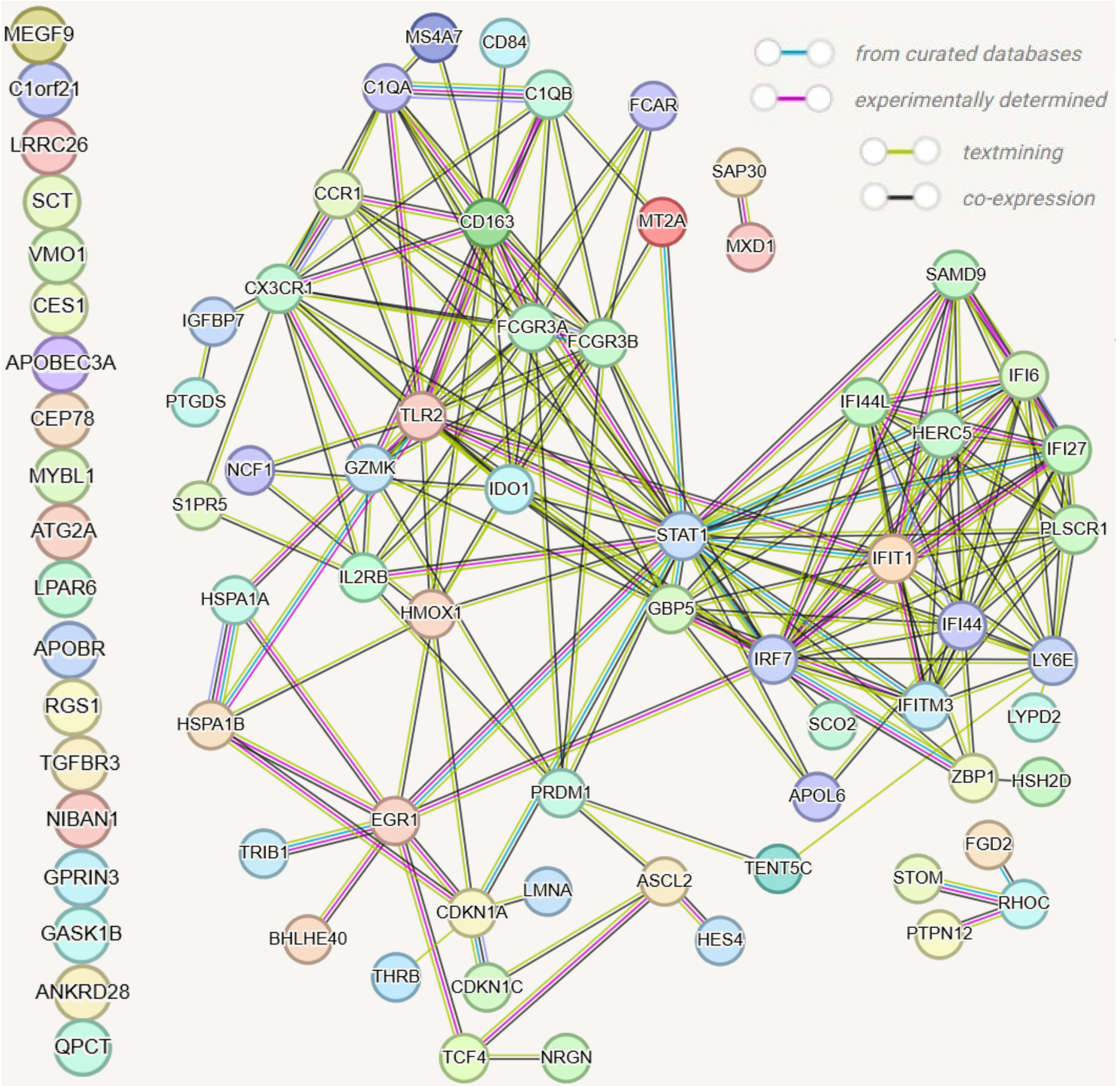


number of nodes: 78
number of edges: 193
average node degree: 4.95
avg. local clustering coefficient: 0.524

expected number of edges: 38
PPI enrichment p-value: < 1.0e-16
*your network has significantly more interactions than expected (what does that mean?)*

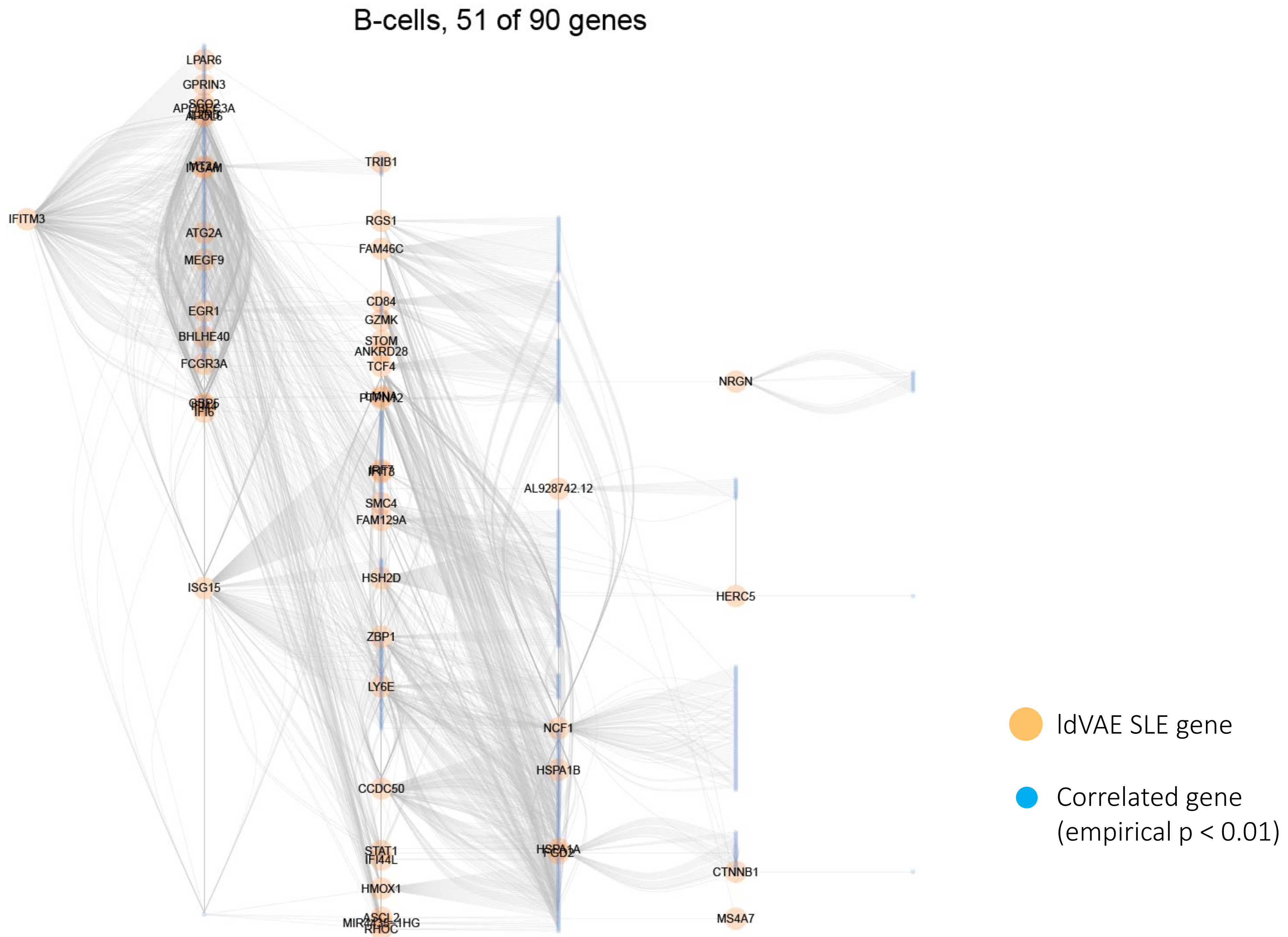


Supplementary Fig. 5A.

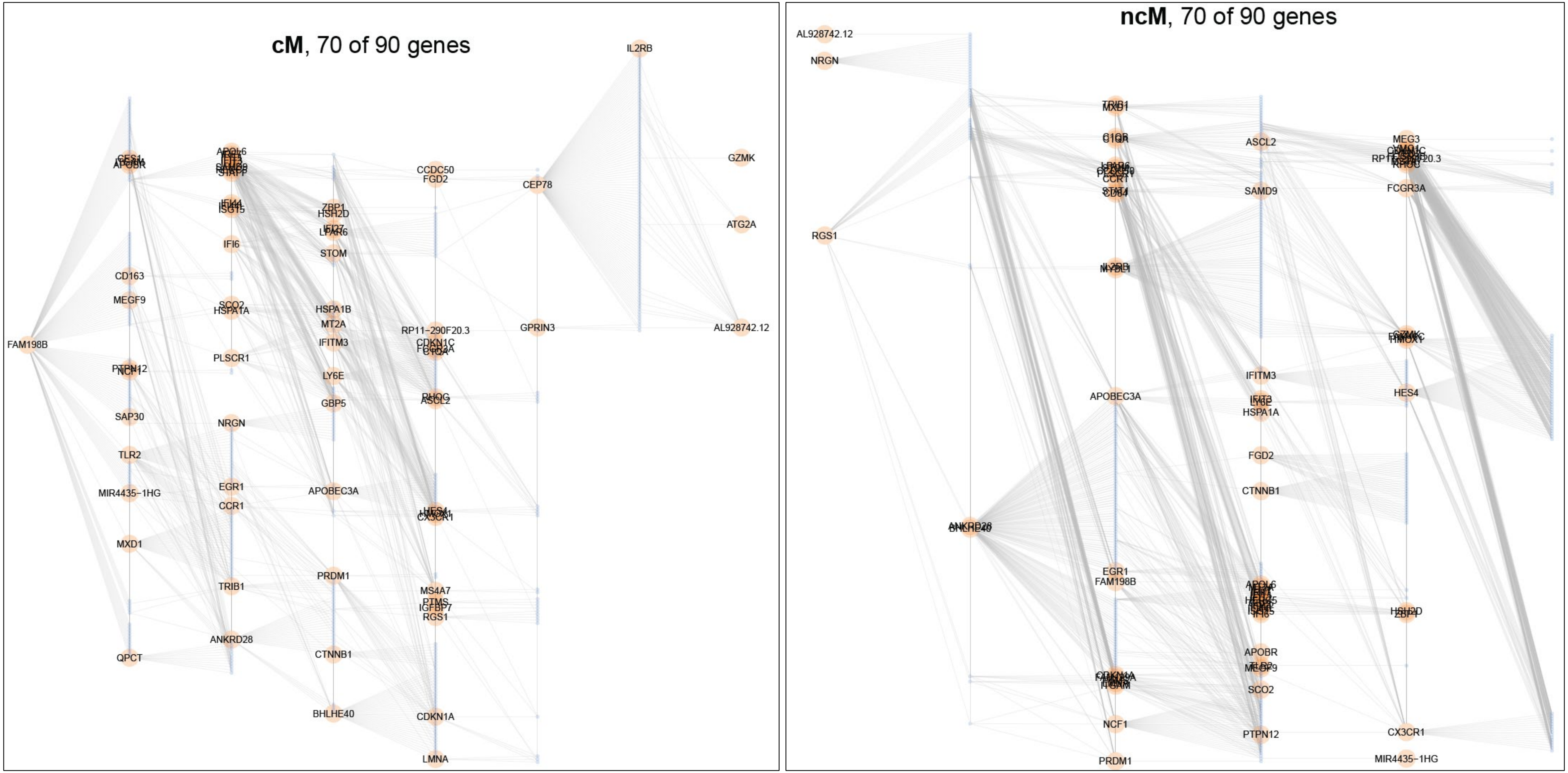


Supplementary Fig. 5B.

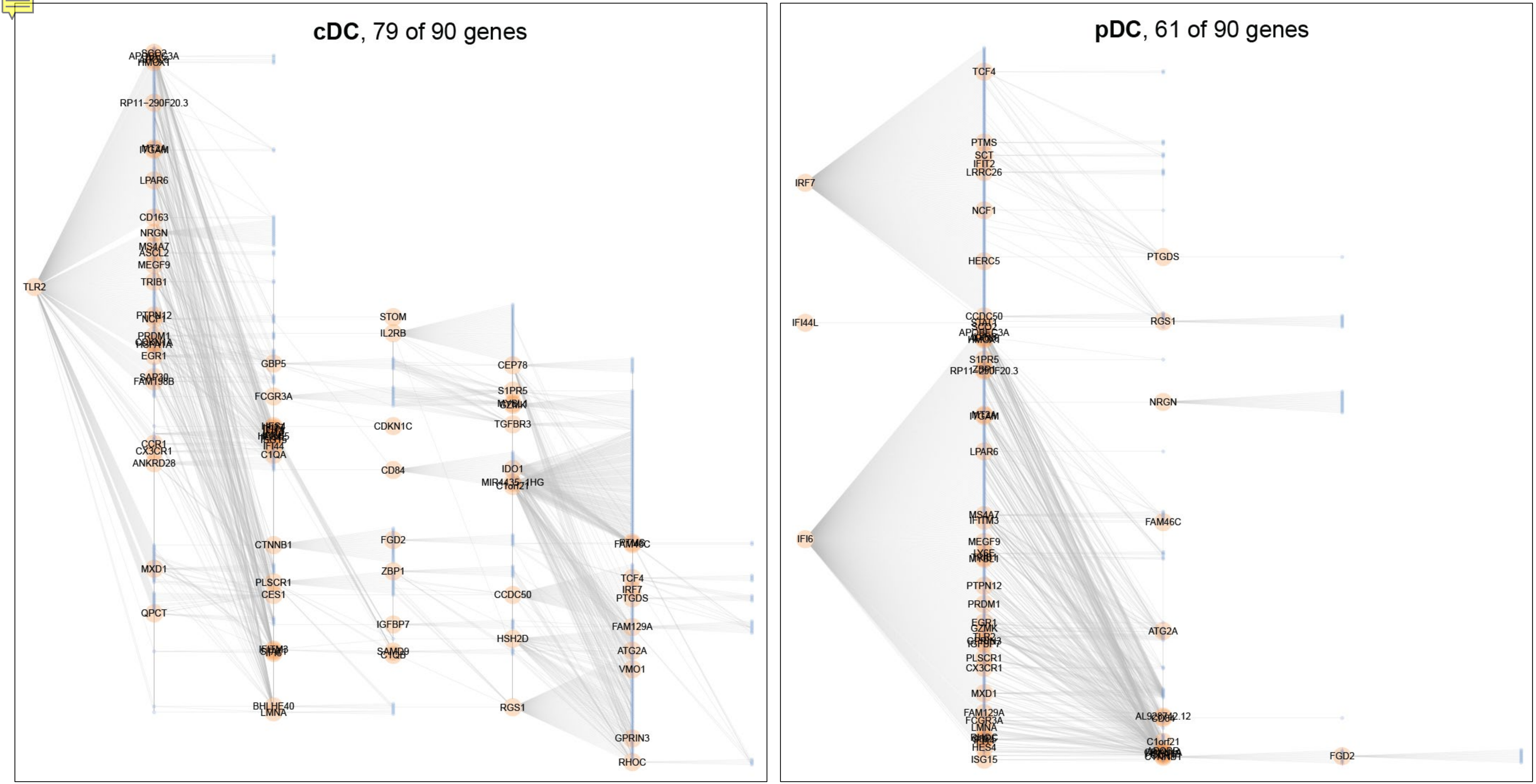


Supplementary Fig. 5C.

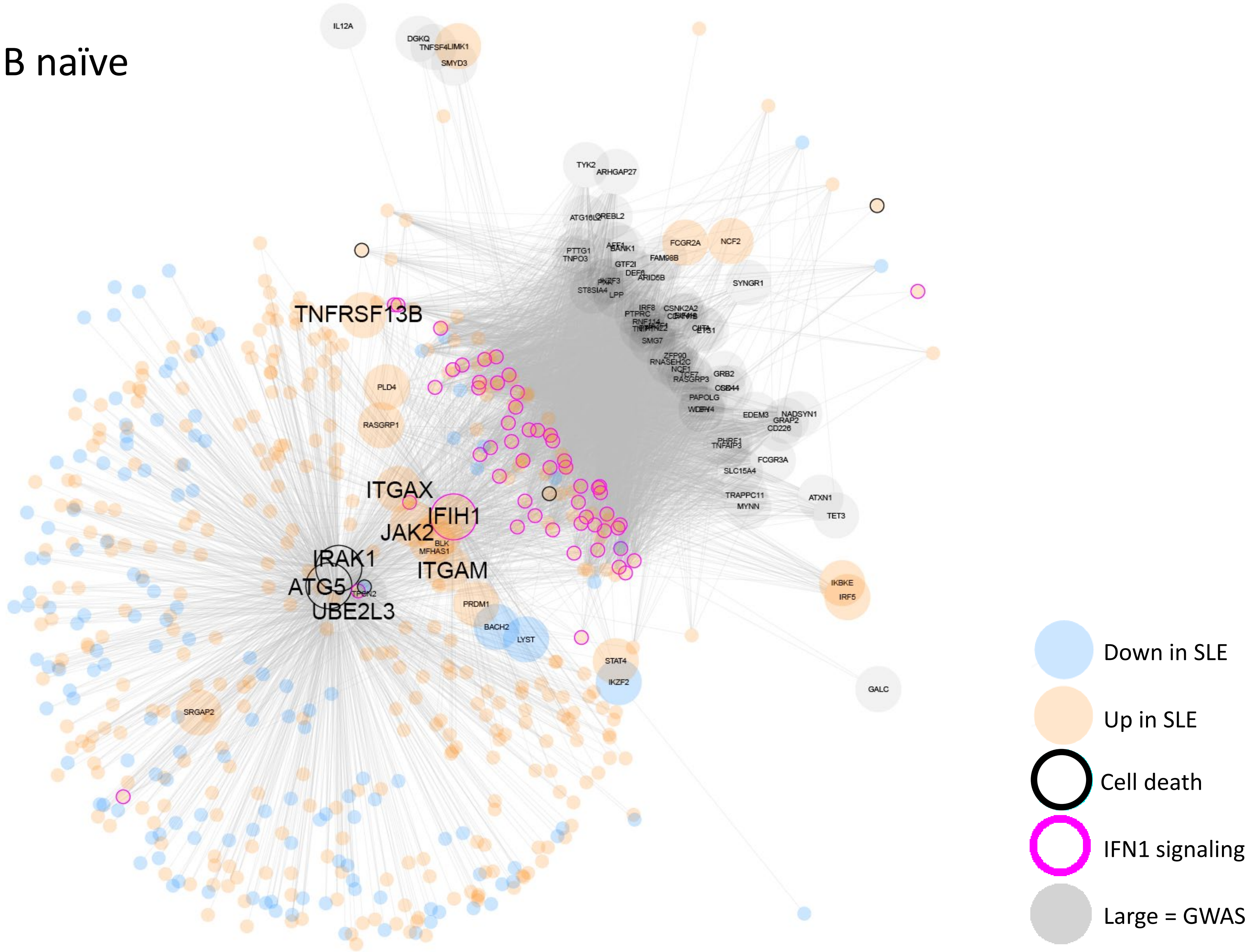


Supplementary Fig. 6A.

### CD16+ Mono

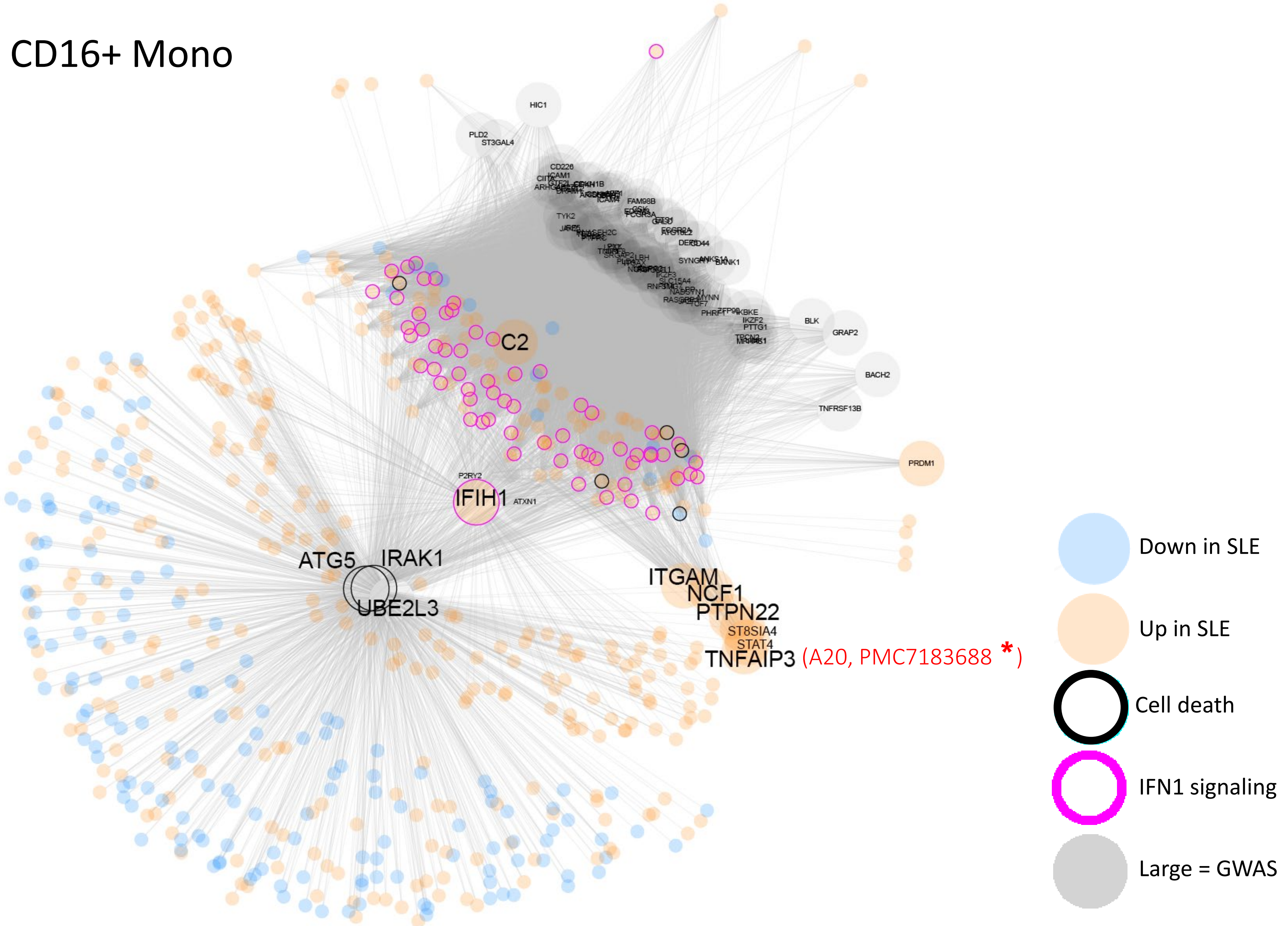


Supplementary Fig. 6B.

### CD14+ Mono

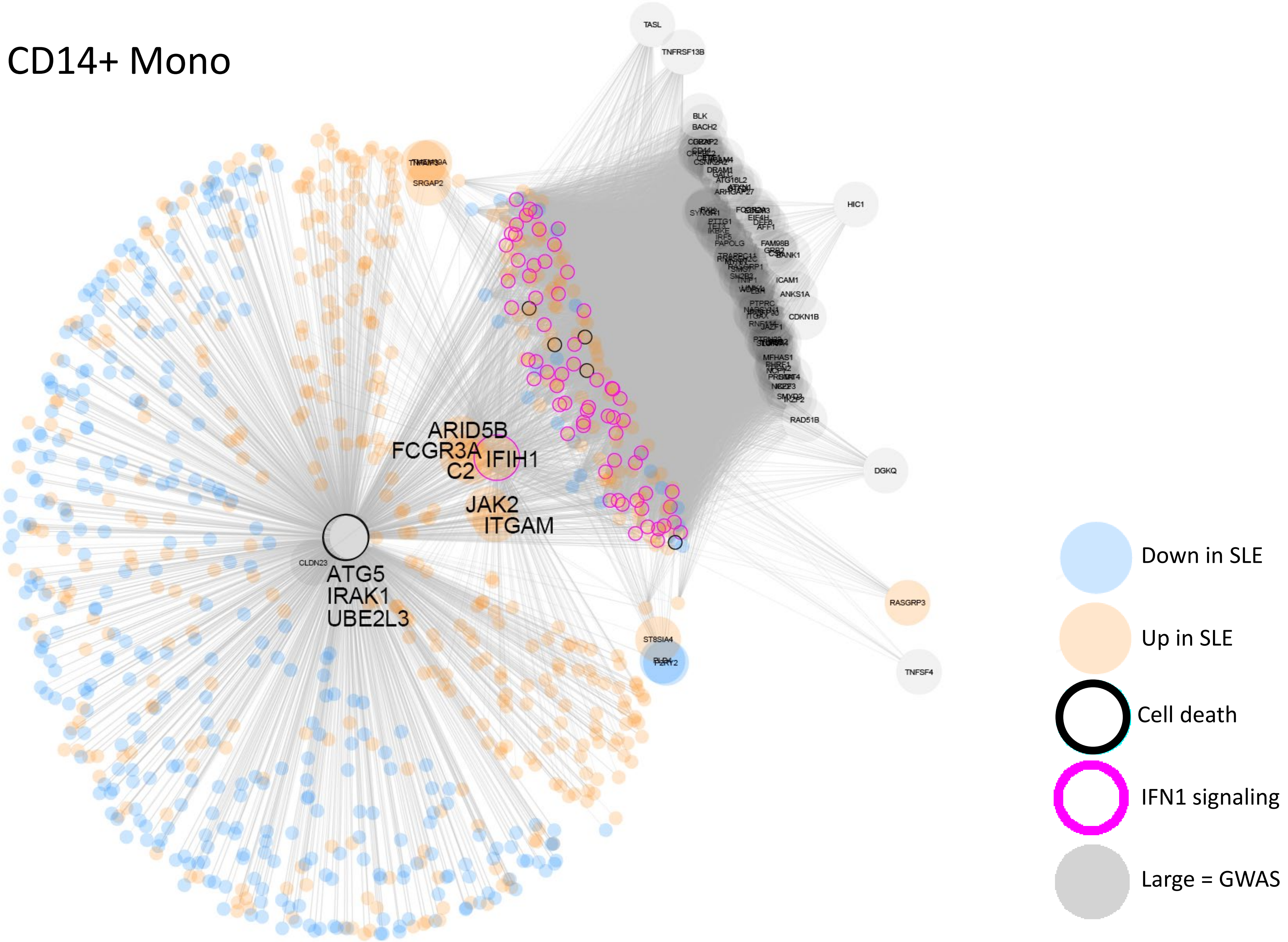


Supplementary Fig. 6C.

pDC

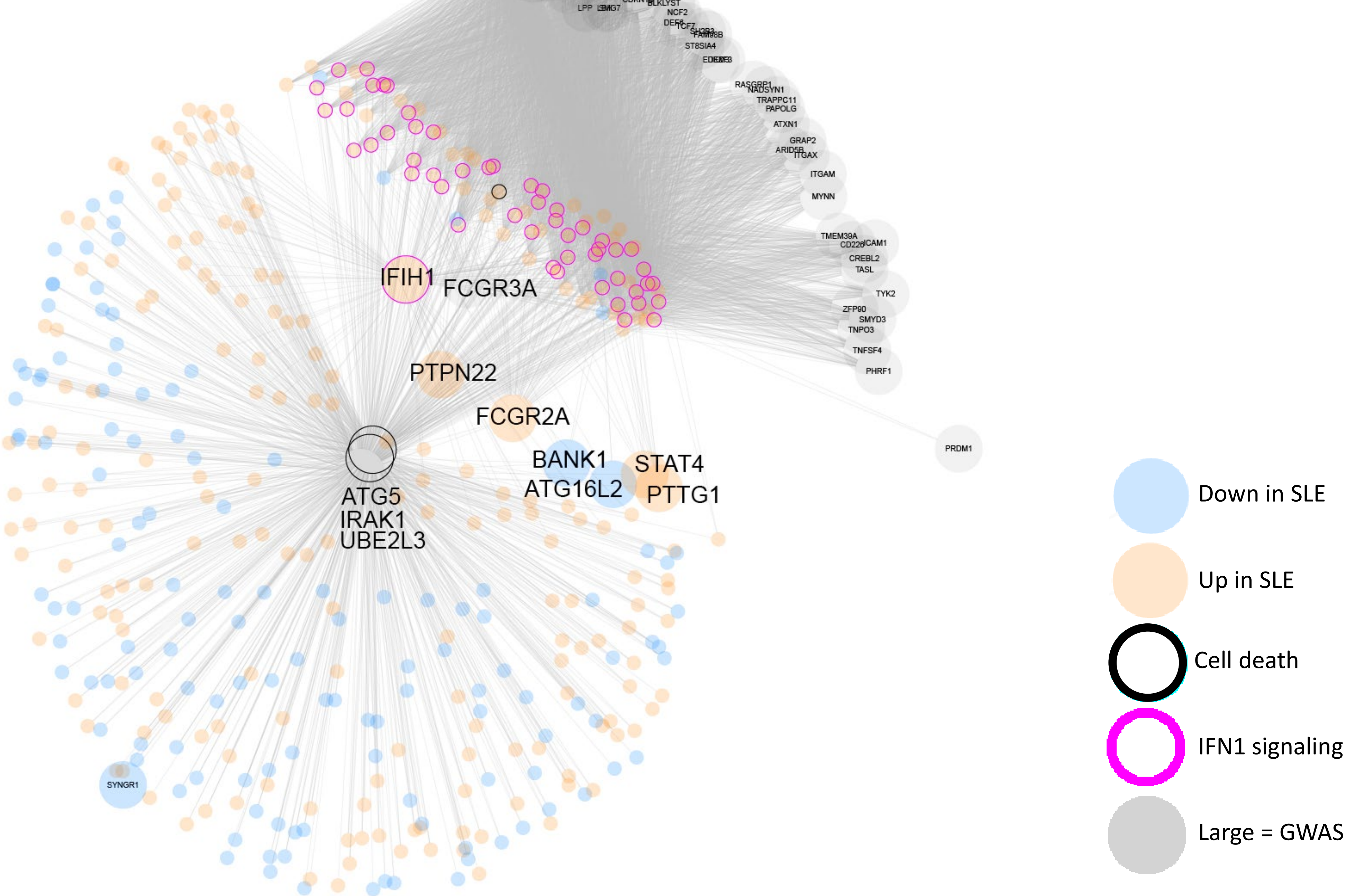


Supplementary Fig. 6D.

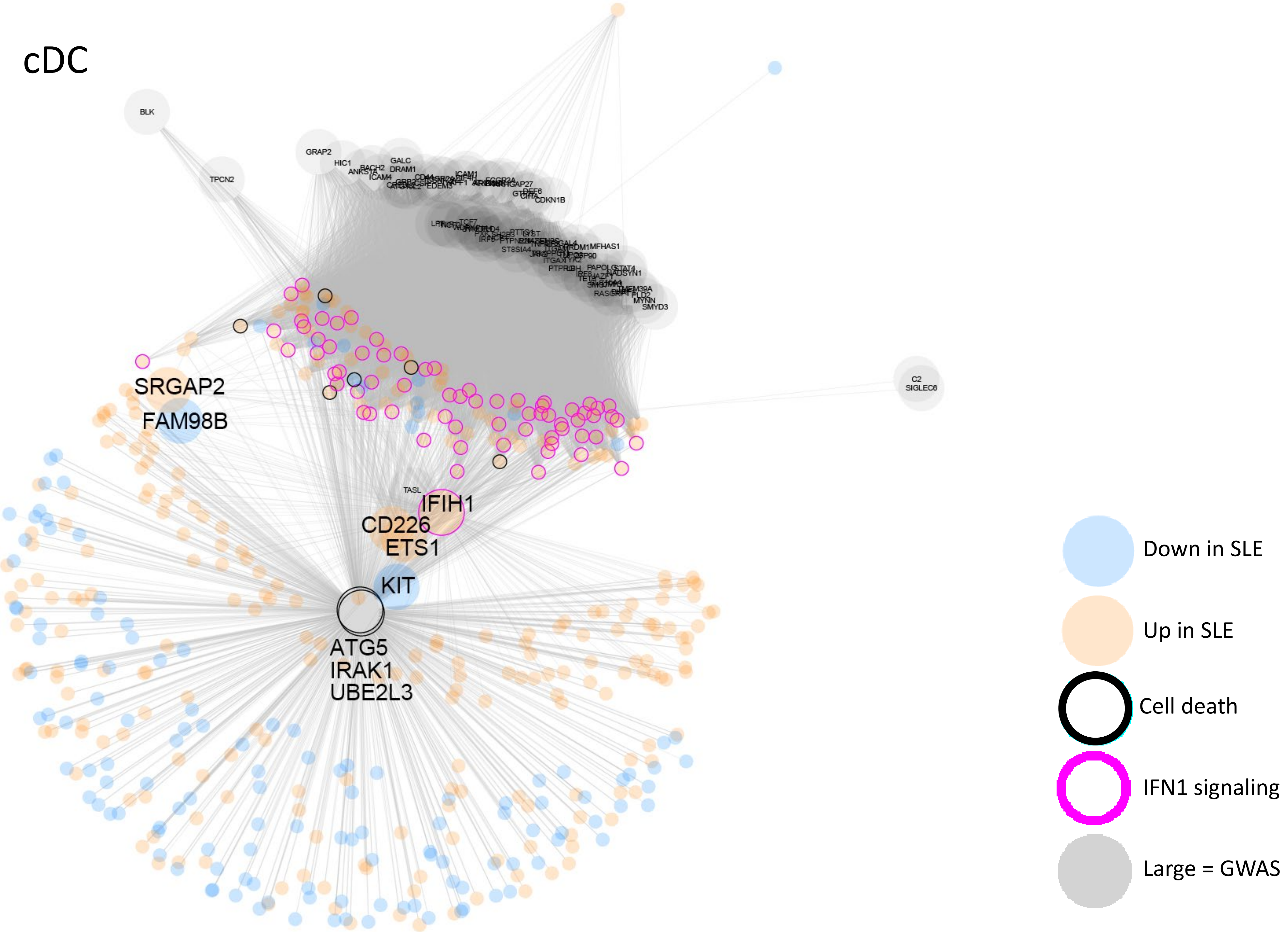


Supplementary Fig. 6E.

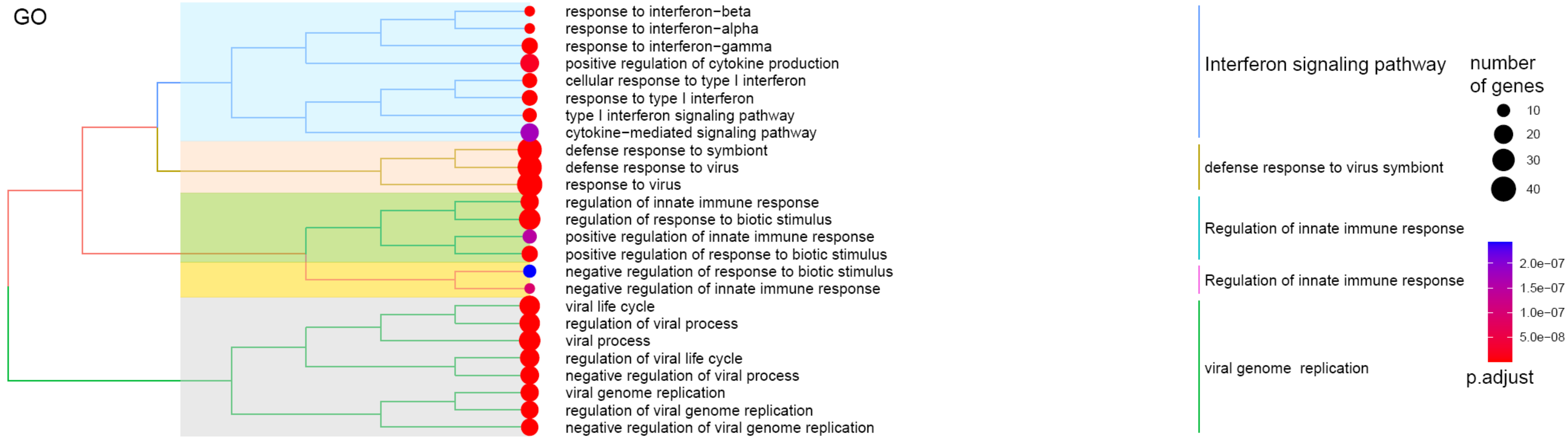


```
  [1] ADAR     ANKAR    APOL6    ARHGAP18 ARHGEF3  ARID5A   B2M      BAX
  [9] BISPR    BLVRB    BRCA2    BST2     BTN3A1   CBR1     CD2AP    CD84
 [17] CHMP5    CMTR1    CYSTM1   DDIT4    DDX58    DDX60    DDX60L   DTX3L
 [25] DYNLT1   EIF2AK2  EPSTI1   EVI2A    FBXO6    FGL2     FKBP5    FOXO1
 [33] GBP1     GBP2     GBP4     GBP5     GRN      HERC5    HIST1H1C HLA-A
 [41] HSH2D    HSPB1    IFI35    IFI44    IFI44L   IFI6     IFIT1    IFIT2
 [49] IFIT3    IFIT5    IFITM1   IFITM2   IFITM3   IRF7     ISG15    ISG20
 [57] ITGAM    ITPR1    JUP      LAP3     LGALS9   LY6E     LY96     MARCKS
 [65] MILR1    MNDA     MT2A     MX1      MX2      NCOA7    NEXN     NLRC5
 [73] NMI      OAS1     OAS2     ODF3B    OPTN     PARP10   PARP11   PARP12
 [81] PARP14   PARP9    PCYOX1   PDE3B    PHF11    PHPT1    PIM1     PLAC8
 [89] PLEKHA1  PLSCR1   PML      PPM1K    PSME2    PVT1     RBM43    RHOB
 [97] RNF213   RPL3     RPL7     RSAD2    SAMD9    SAMD9L   SCCPDH   SESN1
[105] SHISA5   SORL1    SP110    SRGN     STAT1    STAT4    STOM     SYT17
[113] TAP1     TCP11L2  TFEC     TMEM123  TNFSF10  TNFSF13B TRIM22   TRIM5
[121] TSC22D1  TXN      UACA     UBE2L6   VAMP5    VRK2     XAF1     XRN1
[129] ZBP1     ZCCHC2   ZNFX1
```

Supplementary Fig. 7A.

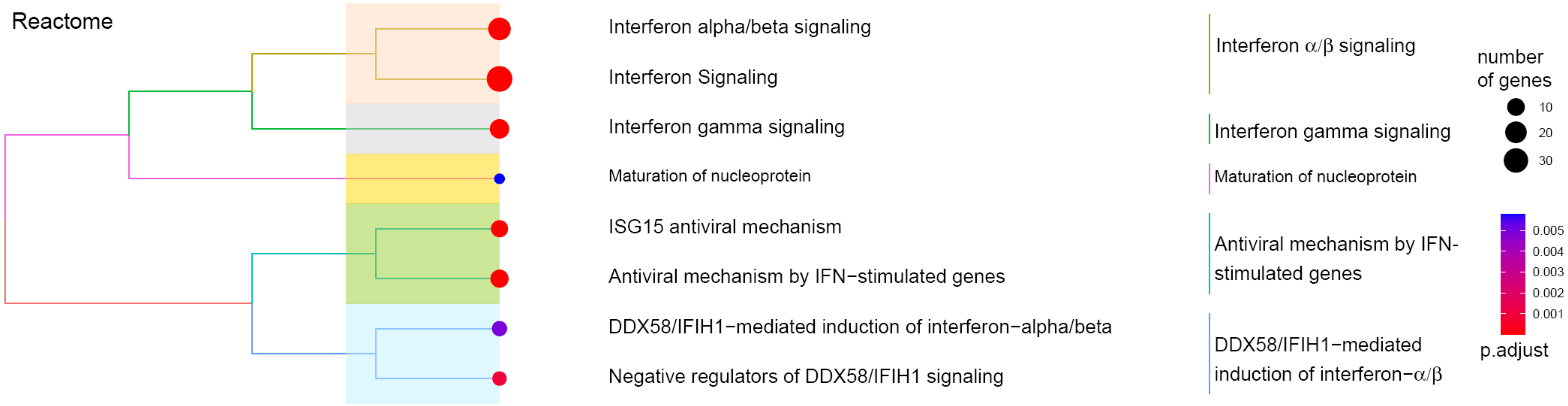


Supplementary Fig. 7B.

### B−naive module

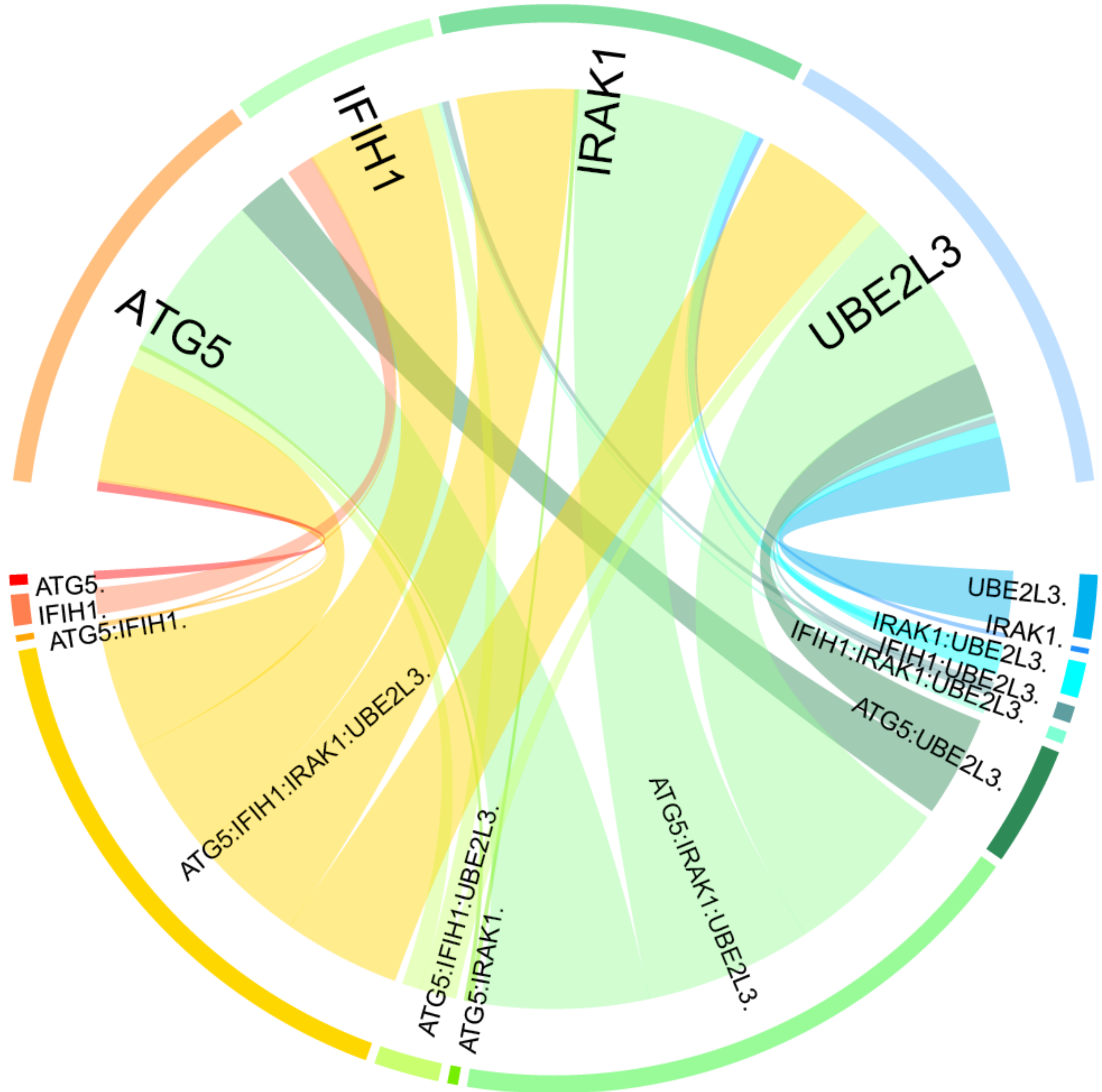


Supplementary Fig. 8A.

CD14 module

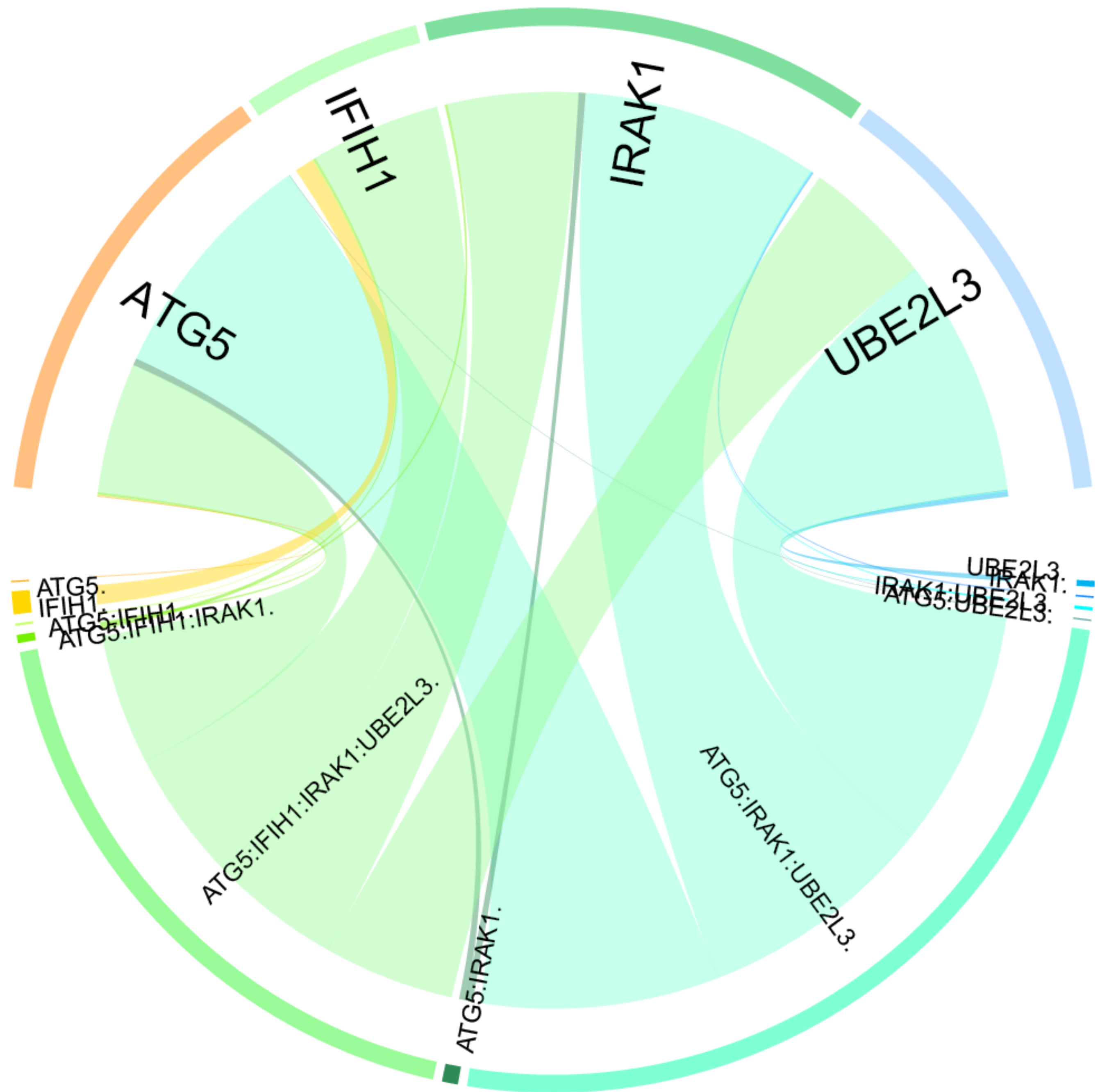


Supplementary Fig. 8B.

### CD16 module

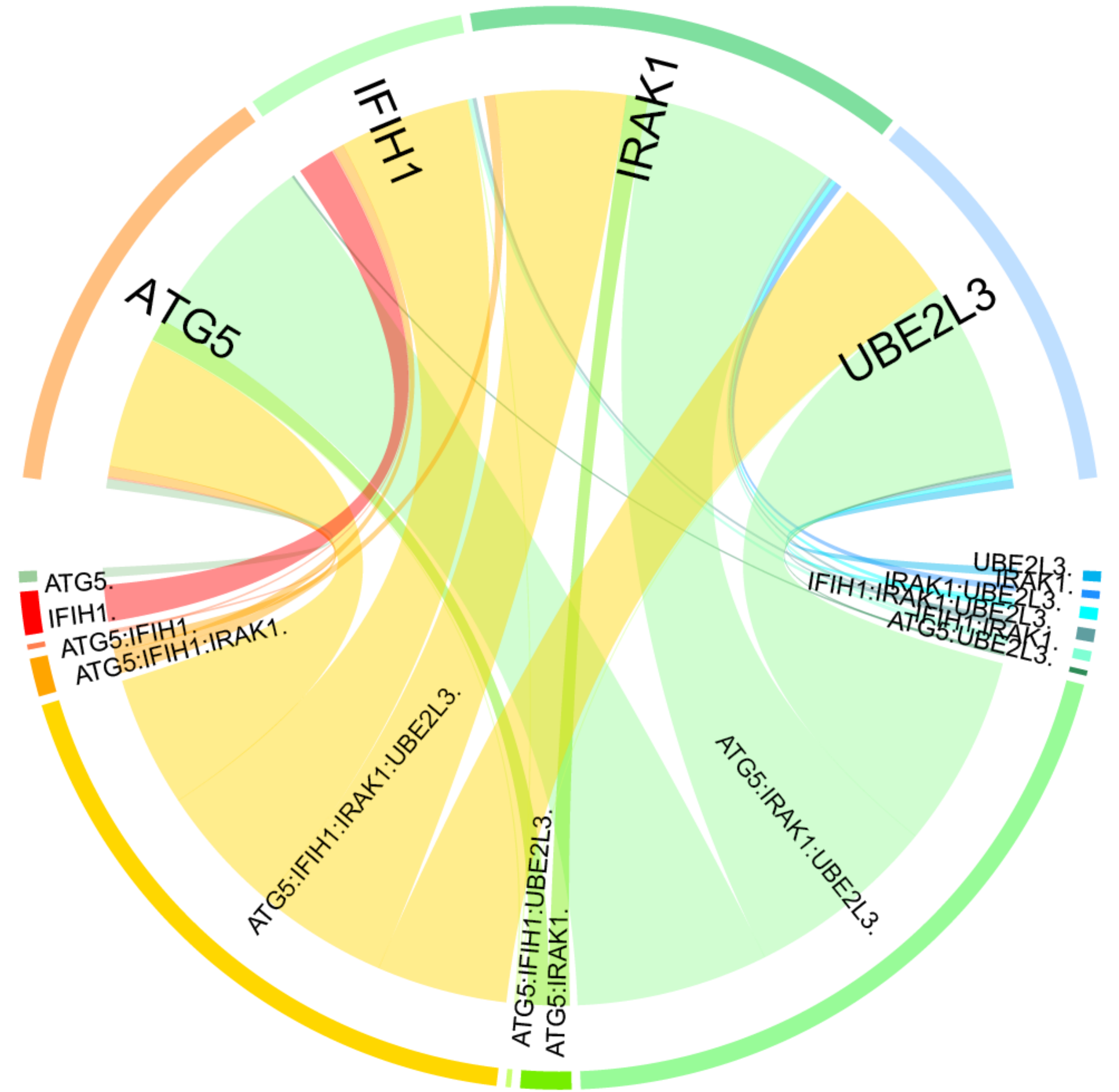


Supplementary Fig. 8C.

### pDC module

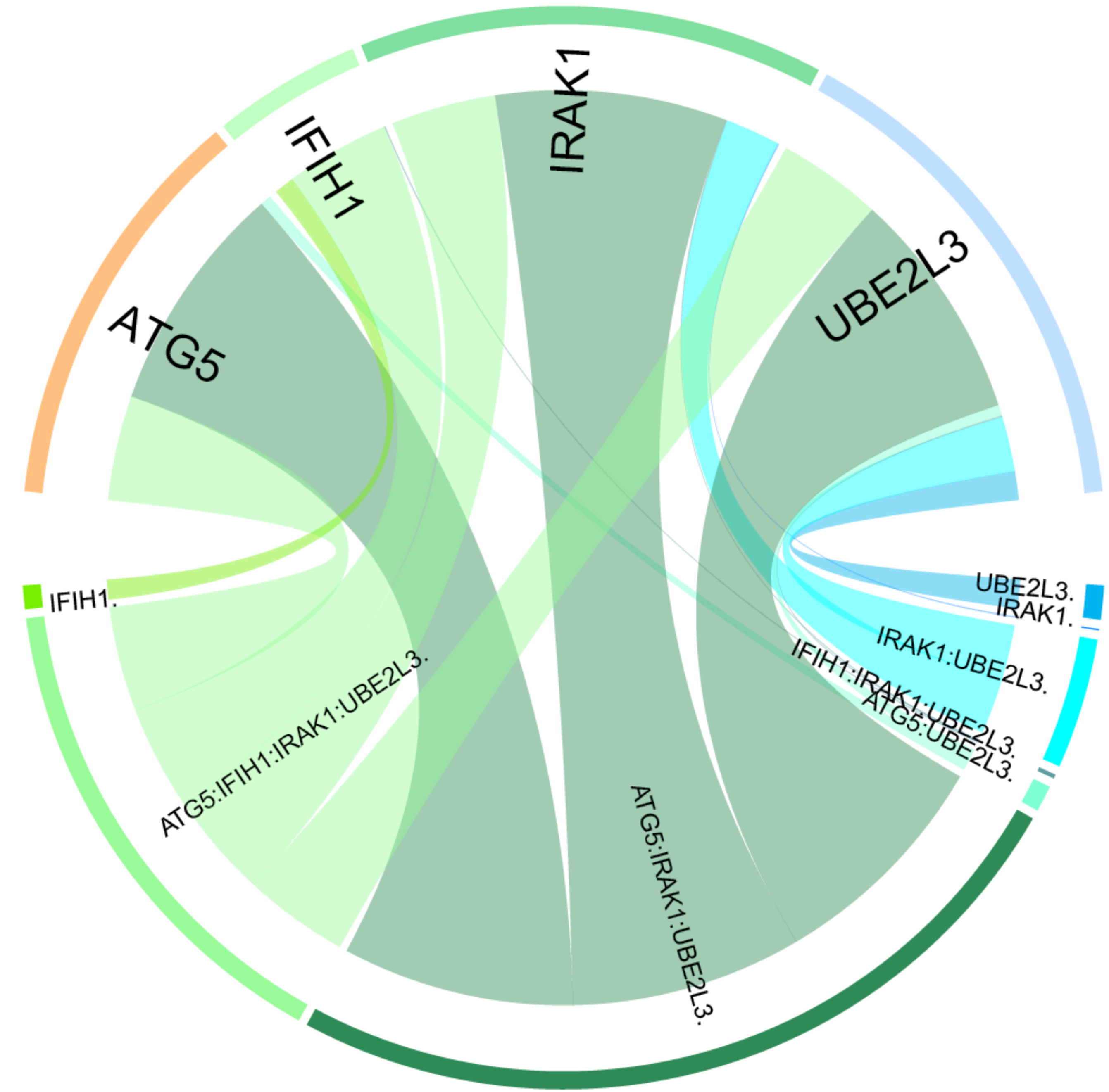


Supplementary Fig. 8D.

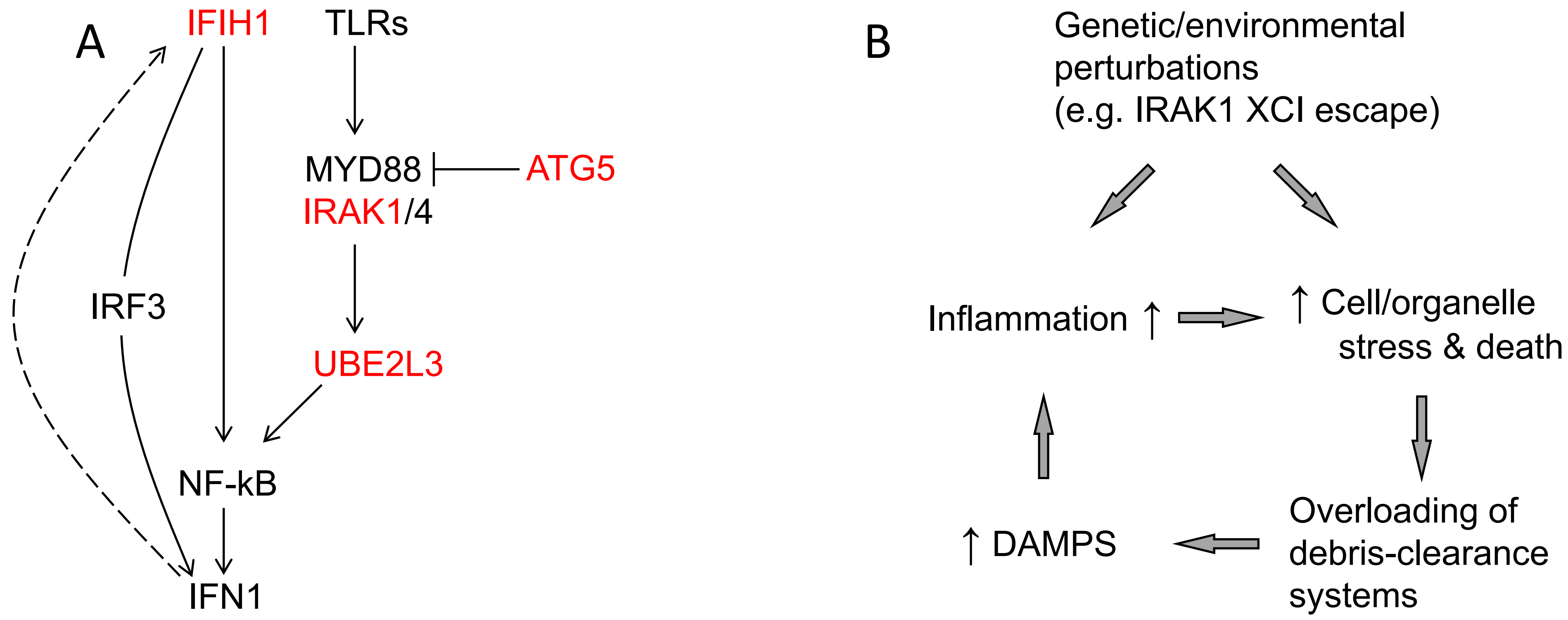


Supplementary Fig. 9.

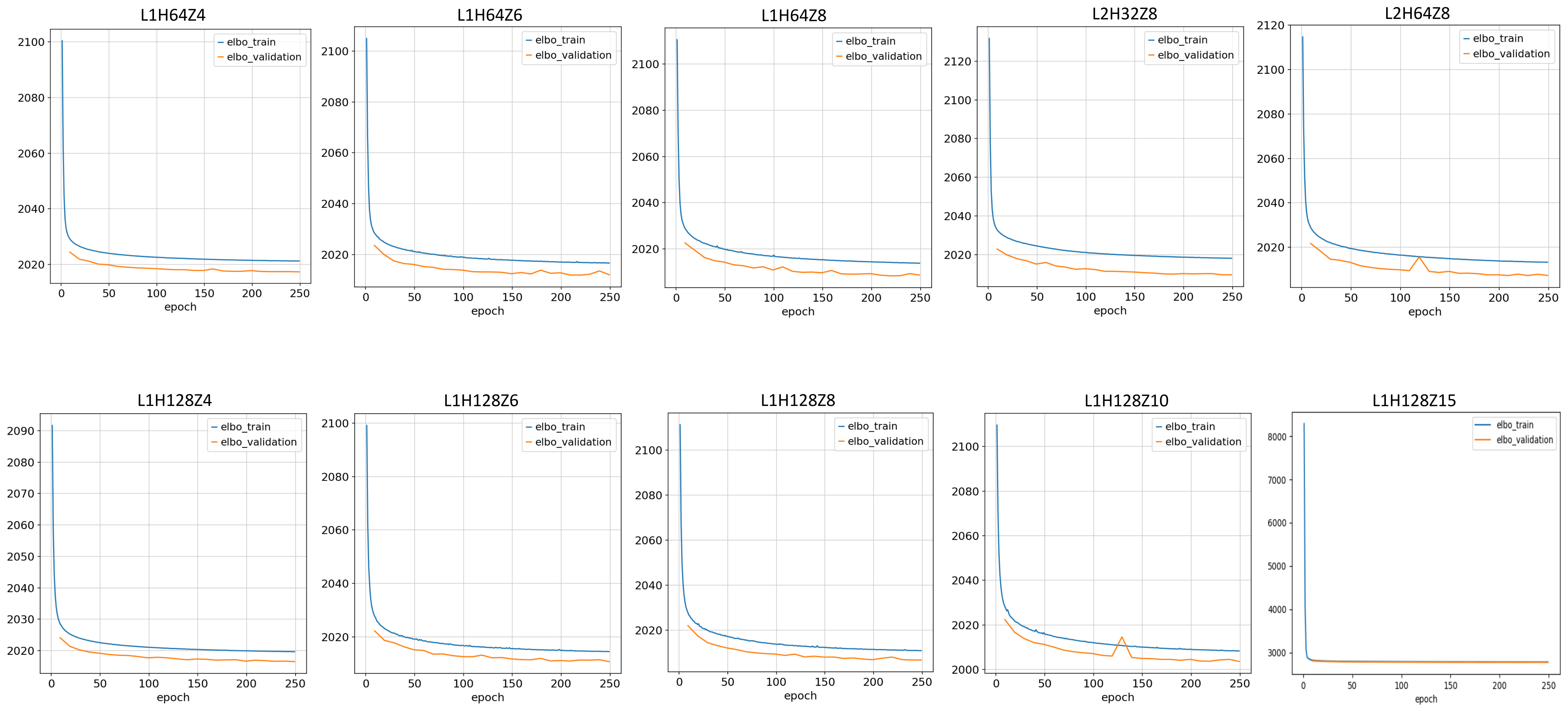


Supplementary Fig. 10.

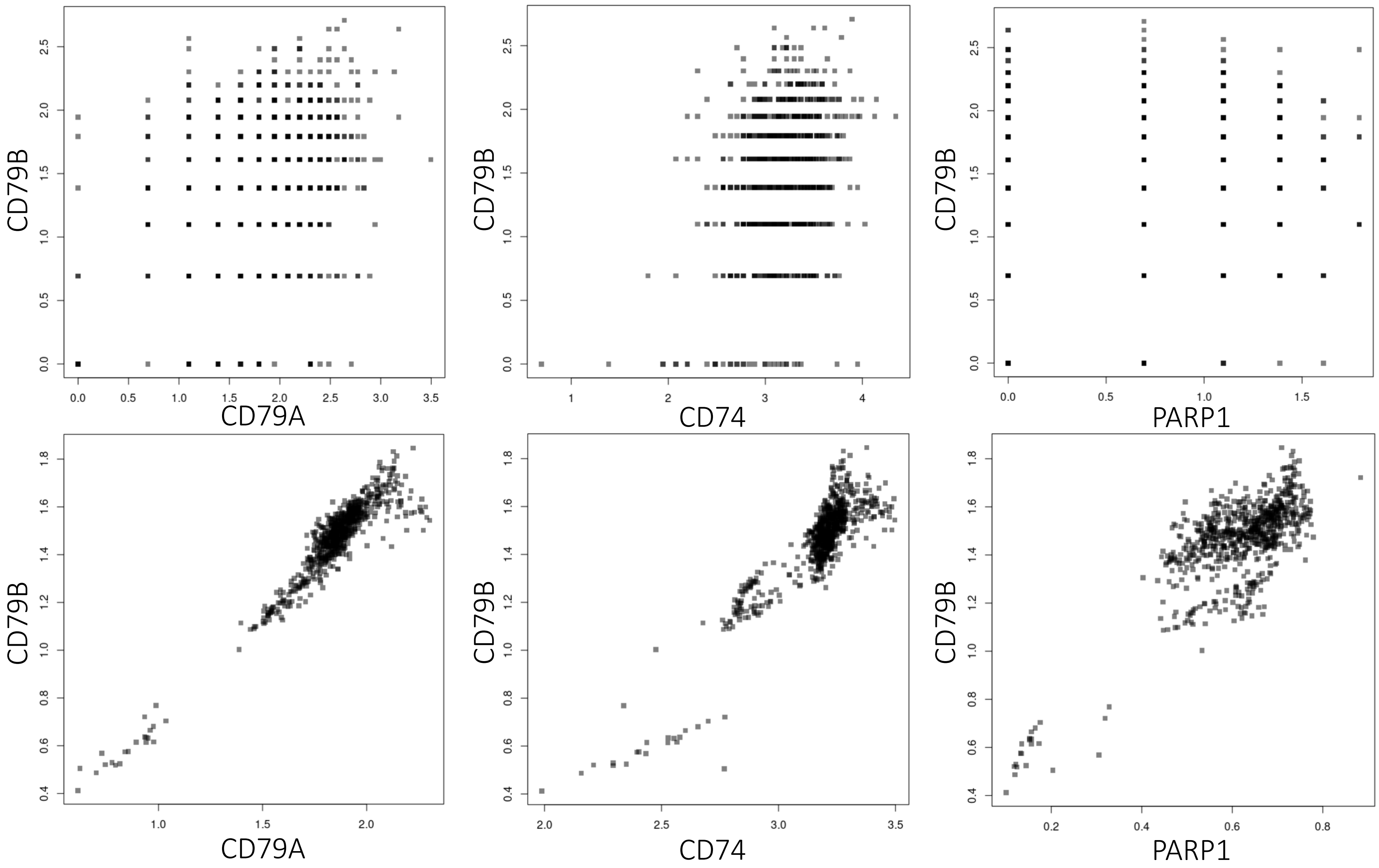


Supplementary Fig. 11.
